# Multi-modal engineering of *Bst* DNA polymerase for thermostability in ultra-fast LAMP reactions

**DOI:** 10.1101/2021.04.15.439918

**Authors:** Inyup Paik, Phuoc H. T. Ngo, Raghav Shroff, Andre C. Maranhao, David J.F. Walker, Sanchita Bhadra, Andrew D. Ellington

## Abstract

DNA polymerase from *Geobacillus stearothermophilus, Bst* DNA polymerase (*Bst* DNAP), is a versatile enzyme with robust strand-displacing activity that enables loop-mediated isothermal amplification (LAMP). Despite its exclusive usage in LAMP assay, its properties remain open to improvement. Here, we describe logical redesign of *Bst* DNAP by using multimodal application of several independent and orthogonal rational engineering methods such as domain addition, supercharging, and machine learning predictions of amino acid substitutions. The resulting Br512g3 enzyme is not only thermostable and extremely robust but it also displays improved reverse transcription activity and the ability to carry out ultrafast LAMP at 74 °C. Our study illustrates a new enzyme engineering strategy as well as contributes a novel engineered strand displacing DNA polymerase of high value to diagnostics and other fields.

## INTRODUCTION

Despite the fact that strand-displacing activity is of great utility for a variety of applications, including isothermal amplification assays, there are relatively few strand-displacing DNA polymerases. In particular, the thermotolerant DNA polymerase from *Geobacillus stearothermophilus (*previously *Bacillus stearothermophilus), Bst* DNA polymerase (*Bst* DNAP), is used in a variety of assays, including loop-mediated isothermal amplification. However, despite its wide use, its properties remain open to improvement, as has been demonstrated by a variety of engineering efforts, including the identification of point mutations that impact its robustness, strand-displacement capabilities, and nascent reverse transcriptase activity.

We hypothesized that in order to greatly improve the core *Bst* DNAP, it would be necessary to improve its thermostability, so that isothermal amplification reactions could be carried out at progressively higher temperatures, which would concomitantly lead to improved, non-enzymatic strand separation and to potentially faster polymerase kinetics. We attempted to improve *Bst* DNAP’s thermostability and functionality through several different, coordinated engineering methods, including the addition of stabilizing domains; the use of machine learning methods to predict stabilizing mutations; and the introduction of charged amino acids that should improve interactions with its polyanionic substrates. By taking multiple engineering tacks, we also were implicitly hypothesizing that changes that focused on different biophysical principles should be orthogonal and additive, as has previously been observed for mutations that impacted different kinetic parameters of an enzyme (1,2). This hypothesis is consistent with findings that enzyme functional improvements are often preceded by general stabilizing mutations (3).

Fusion domains have previously been used in the construction of thermostable DNA polymerases with improved properties for PCR, as opposed to isothermal amplification. In the case of Phusion DNA polymerase, the addition of the Sso7d gene, a DNA binding protein from *Sulfolobus solfactaricus*, stabilizes the polymerase/DNA complex and enhances the processivity by up to 9 times, allowing longer amplicons in less time with less influence of PCR inhibitors (4). In order to both increase the thermostability of the *Bst* DNAP and to provide it with greater interactions with its polyanionic DNA substrate, we similarly identified an extremely robust protein fusion partner based on the villin headpiece (5). The terminal thirty-five amino acids of the headpiece (HP35) consists of three α-helices that form a highly conserved hydrophobic core (6), which exhibits co-translational, ultrafast, and autonomous folding properties that may circumvent kinetic traps during protein folding (7). The ultrafast folding property of the villin headpiece subdomain has made it a model for protein folding dynamics and simulation studies (8). In addition, HP35 displays thermostability with a transition midpoint (Tm) of 70°C (6,7).

## MATERIAL AND METHODS

### Chemicals and reagents

All chemicals were of analytical grade and were purchased from Sigma-Aldrich (St. Louis, MO, USA) unless otherwise indicated. All commercially sourced enzymes and related buffers were purchased from New England Biolabs (NEB, Ipswich, MA, USA) unless otherwise indicated. All oligonucleotides and gene blocks (Supplementary Table 1) were obtained from Integrated DNA Technologies (IDT, Coralville, IA, USA). SARS-CoV-2 genomic RNA and inactivated virions were obtained from American Type Culture Collection, Manassas, VA, USA.

### Br512 and enzyme variants purification protocol

The overall scheme for Br512 and its variants purification is shown in Supplementary Figure 20. In short, Br512 was cloned into an in-house E. coli expression vector under the control of a T7 RNA polymerase promoter (pKAR2). Full sequence and annotations of the pKAR2-Br512 plasmid are available in Supplementary Table 2. The Br512 expression construct pKAR2-Br512 and its variants were then transformed into E. coli BL21(DE3) (NEB, C2527H). A single colony was seed cultured overnight in 5 mL of superior broth (Athena Enzyme Systems, 0105). The next day, 1 mL of seed culture was inoculated into 1 L of superior broth and grown at 37 OC until it reached an OD600 of 0.5-0.6. Enzyme expression was induced with 1 mM IPTG and 100 ng/mL of anhydrous tetracycline (aTc) at 18 OC for 18 h (or overnight). The induced cells were pelleted at 5000xg for 10 min at 4OC and resuspended in 30mL of ice-cold lysis buffer (50 mM Phosphate Buffer, pH 7.5, 300 mM NaCl, 20 mM imidazole, 0.1% Igepal CO-630, 5 mM MgSO4, 1 mg/mL HEW Lysozyme, 1x EDTA-free protease inhibitor tablet, Thermo Scientific, A32965). The samples were then sonicated (1 sec ON, 4 sec OFF) for a total time of 4 minutes with 40% amplitude. The lysate was centrifuged at 35,000 xg for 30 minutes at 4 °C. The supernatant was transferred to a clean tube and filtered through a 0.2 μm filter.

Protein from the supernatant was purified using metal affinity chromatography on a Ni-NTA column. Briefly, 1mL of Ni-NTA agarose slurry was packed into a 10mL disposable column and equilibrated with 20 column volume (CV) of equilibration buffer (50 mM Phosphate Buffer, pH 7.5, 300 mM NaCl, 20 mM imidazole). The sample lysate was loaded onto the column and the column was developed by gravity flow. Following loading, the column was washed with 20 CV of equilibration buffer and 5 CV of wash buffer (50 mM Phosphate Buffer, pH 7.5, 300 mM NaCl, 50 mM imidazole). Br512 was eluted with 5 mL of elution buffer (50 mM Phosphate Buffer, pH 7.5, 300 mM NaCl, 250 mM imidazole). The eluate was dialyzed twice with 2L of Ni-NTA dialysis buffer (40 mM Tris-HCl, pH 7.5, 100 mM NaCl, 1 mM DTT, 0.1% Igepal CO-630). The dialyzed eluate was further passed through an equilibrated 5 mL heparin column (HiTrap^™^ Heparin HP) on a FPLC (AKTA pure, GE healthcare) and eluted using a linear NaCl gradient generated from heparin buffers A and B (40 mM Tris-HCl, pH 7.5, 100 mM NaCl for buffer A; 2M NaCl for buffer B, 0.1% Igepal CO-630). The collected final eluate was dialyzed first with 2 L of heparin dialysis buffer (50 mM Tris-HCl, pH 8.0, 50 mM KCl, 0.1% Tween-20) and second with 2 L of final dialysis buffer (50% Glycerol, 50 mM Tris-HCl, pH 8.0, 50 mM KCl, 0.1% Tween-20, 0.1% Igepal CO-630, 1 mM DTT). The purified Br512 was quantified by Bradford assay and SDS-PAGE/coomassie gel staining alongside a bovine serum albumin (BSA) standard.

### Real-time *GAPDH* LAMP-OSD

LAMP-OSD reaction mixtures were prepared in 25 μL volume containing indicated amounts of human glyceraldehyde-3-phosphate dehydrogenase (*GAPDH*) DNA templates along with a final concentration of 1.6 μM each of BIP and FIP primers, 0.4 μM each of B3 and F3 primers, and 0.8 μM of the loop primer. Amplification was performed in one of the following buffers – 1X Isothermal buffer (NEB) (20 mM Tris-HCl, 10 mM (NH4)2SO4, 50 mM KCl, 2 mM MgSO4, 0.1% Tween 20, pH 8.8 at 25 °C), G6B buffer (60 mM Tris-HCl, pH 8.0, 2 mM (NH4)2SO4, 40 mM KCl, 4 mM MgCl2), G1B buffer (60 mM Tris-HCl, pH 8.0, 5 mM (NH4)2SO4, 10 mM KCl, 4 mM MgSO4, 0.01% Triton X-100), G2A buffer (20 mM Tris-HCl, pH 8.0, 5 mM (NH4)2SO4, 10 mM KCl, 4 mM MgCl2, 0.01% Triton X-100), or G3A buffer (60 mM Tris-HCl, pH 8.0, 10 mM KCl, 4 mM MgCl2, 0.01% Triton X-100). The buffer was appended with 1 M betaine, 0.4 mM dNTPs, 2 mM additional MgSO4 (only for reactions in Isothermal buffer), and either Bst 2.0 DNA polymerase (16 units), Bst-LF DNA polymerase (20 pm), or Br512 DNA polymerase (0.2 pm, 2 pm, 20 pm, or 200 pm). Assays read using OSD probes received 100 nM of OSD reporter prepared by annealing 100 nM fluorophore-labeled OSD strands with a 5-fold excess of the quencher-labeled OSD strands by incubation at 95 °C for 1 min followed by cooling at the rate of 0.1 °C/sec to 25 °C. Assays read using intercalating dyes received 1X EvaGreen (Biotium, Freemont, CA, USA) instead of OSD probes. For real-time signal measurement these LAMP reactions were transferred into a 96-well PCR plate, which was incubated in a LightCycler 96 real-time PCR machine (Roche, Basel, Switzerland) maintained at 65 °C for 90 min. Fluorescence signals were recorded every 3 min in the FAM channel and analyzed using the LightCycler 96 software. For assays read using EvaGreen, amplification was followed by a melt curve analysis on the LightCycler 96 to distinguish target amplicons from spurious background.

### Endpoint *GAPDH* LAMP-OSD

LAMP-OSD reaction mixtures for visual endpoint readout of OSD fluorescence were prepared in 25 μL volume containing either 1X Isothermal buffer and 16 units of Bst 2.0 or G6B buffer and 20 pm of Br512. The reactions also contained 1 M betaine, 1.4 mM dNTPs, 2 mM additional MgSO4 (only for reactions in Isothermal buffer), and 100 nM OSD reporter prepared by annealing 100 nM fluorophore-labeled OSD strands with a 2-fold excess of the quencher-labeled OSD strands by incubation at 95 °C for 1 min followed by cooling at the rate of 0.1 °C/sec to 25 °C. A final concentration of 1.6 μM each of BIP and FIP primers, 0.4 μM each of B3 and F3 primers, and 0.8 μM of the loop primer were added to some reactions while control assays without primers received the same volume of water. Some assays were seeded with 3 μL of human saliva that had been heated at 95 °C for 10 min while other assays received the same volume of water. All assays were incubated in a thermocycler maintained at 65 °C for 60 min following which endpoint OSD fluorescence was imaged using a ChemiDoc camera (Bio-Rad Laboratories, Hercules, CA, USA).

### SARS-CoV-2 RT-LAMP-OSD assays

Individual 25 μL RT-LAMP-OSD assays were assembled either in 1X Isothermal buffer containing 16 units of Bst 2.0 or in G6D buffer (60 mM Tris-HCl, pH 8.0, 2 mm (NH4)2SO4, 40 mM KCl, 8 mM MgCl2) containing 20 pm of Br512, 20 pm of Bst-LF, or 16 units of Bst 2.0. The buffer was supplemented with 1.4 mM dNTPs, 0.4 M betaine, 6 mM additional MgSO4 (only for reactions in Isothermal buffer), and 2.4 μM each of FIP and BIP, 1.2 μM of indicated loop primers, and 0.6 μM each of F3 and B3 primers. Amplicon accumulation was measured by adding OSD probes. First, Tholoth, Lamb, and NB OSD probes were prepared by annealing 1 μM of the fluorophore-labeled OSD strand with 2 μM, 3 μM, and 5 μM, respectively of the quencher-labeled strand in 1X Isothermal buffer. Annealing was performed by denaturing the oligonucleotide mix at 95 °C for 1 min followed by slow cooling at the rate of 0.1 °C/s to 25 °C. Excess annealed probes were stored at −20 °C. Annealed Tholoth, Lamb, and NB OSD probes were added to their respective RT-LAMP reactions at a final concentration of 100 nM of the fluorophore-bearing strand. The assays were seeded with indicated amounts of SARS-CoV-2 viral genomic RNA in TE buffer (10 mM Tris-HCl, pH 7.5, 0.1 mM EDTA) and either incubated for 1 h in a thermocycler maintained at 65 °C for endpoint readout or transferred to a 96-well plate and incubated in the LightCycler 96 real-time PCR machine maintained at 65 °C for real time measurement of amplification kinetics. Endpoint OSD fluorescence was read visually using a blue LED torch with orange filter and imaged using a cellphone camera or a ChemiDoc (Bio-Rad) camera. OSD fluorescence in assays incubated in the real-time PCR machine was measured every 3 min in the FAM channel and analyzed using the LightCycler 96 software.

Multiplex RT-LAMP-OSD assays comprising 6-Lamb and NB primers and OSD probes were set up using the same conditions as above except that the total LAMP primer amounts were made up of equimolar amounts of 6-Lamb and NB primers supplemented with 0.2 μM each of additional NB FIP and BIP primers. Some multiplex assays received 30 pm Br512 instead of 20 pm enzyme. Some Br512 multiplex assays also received 20 units of the RNase inhibitor, SUPERase.In (Thermo Fisher Scientific, Waltham, MA). Multiplex assays were seeded with indicated amounts of either SARS-CoV-2 viral genomic RNA or inactivated virions in the presence of either 3 μL of TE buffer or human saliva pre-heated at 95 °C for 10 min. The assays were either incubated for 1 h in a thermocycler maintained at 65 °C for endpoint readout or transferred to a 96-well plate and incubated in the LightCycler 96 real-time PCR machine maintained at 65 °C for real time measurement of amplification kinetics. Endpoint OSD fluorescence was read visually using a blue LED torch with orange filter and imaged using a cellphone camera or a ChemiDoc (Bio-Rad) camera. OSD fluorescence in assays incubated in the real-time PCR machine was measured every 3 min in the FAM channel and analyzed using the LightCycler 96 software.

### Lyophilization of Br512

Multiplex SARS-CoV-2 assay reagent mixes were prepared by combining dNTPs, NB and 6-Lamb primers and OSD probes, trehalose, and glycerol-free Br512 in the following amounts per individual reaction – (i) 35 nanomoles of dNTPs, (ii) 30 pm each of 6-Lamb FIP and BIP, 15 pm each of 6-Lamb LF and LB loop primers, 7.5 pm each of 6-Lamb F3 and B3 primers, (iii) 35 pm each of NB FIP and BIP, 15 pm of NB LB loop primer, 7.5 pm each of NB F3 and B3 primers, (iv) 2.5 pm of NB fluorophore labeled OSD strand pre-annealed with 5-fold excess of the quencher labeled OSD strand, (v) 2.5 pm of Lamb fluorophore labeled OSD strand pre-annealed with a 3-fold excess of the quencher labeled OSD strand, (vi) 1.25 micromoles of trehalose, and (vii) 20 pm or 30 pm of BR512. The reagent mixes were distributed in 0.2 mL PCR tubes and frozen for 1 h on dry ice prior to lyophilization for 2.5 h at 197 mTorr and −108 °C using the automated settings in a VirTis Benchtop Pro lyophilizer (SP Scientific, Warminster, PA, USA). Lyophilized assays were stored with desiccant at −20 °C until use.

Lyophilized assays were rehydrated immediately prior to use by adding 22 μL of 1X G6D buffer containing 10 micromoles of betaine. Rehydrated assays were seeded with indicated amounts of SARS-CoV-2 viral genomic RNA in a total volume of 3 μL and incubated for 1 h in a thermocycler maintained at 65 °C. Endpoint OSD fluorescence was read visually using a blue LED torch with orange filter and imaged using a cellphone camera.

### Site directed mutagenesis

Site directed mutagenesis was performed using Q5^®^ Site-Directed Mutagenesis Kit from NEB (E0554S) according to manufacturer’s instructions. The pKAR2-Br512 plasmid was used as a template to introduce mutations suggested by the Mutcompute and Supercharging analysis. All primer sequences are listed in Supplementary Table 1. The introduced mutations on the plasmids were confirmed by Sanger sequencing.

#### Heat challenge and high temperature LAMP

LAMP reaction mixtures were prepared in 25μL volume containing 10pg of human glyceraldehyde-3-phosphate dehydrogenase (*GAPDH*) DNA template plasmid along with a final concentration of 1.6 μM each of BIP and FIP primers, 0.4 μM each of B3 and F3 primers, and 0.8 μM of the loop primer. The reaction mixtures were preassemble on ice and aliquoted into PCR tubes. A total 20pm of enzyme variants were added to the wells. Amplification was performed in the following buffer (1X LAMP heat challenge buffer: 40 mM Tris-HCl, pH 8.0, 10 mM (NH4)2SO4, 80 mM KCl, 4mM MgCl2) supplemented with 0.4mM dNTP, 1X Evagreen Dye, and 0.4M betaine unless otherwise indicated. For heat challenges, PCR tubes that contain the reaction mixtures and 20pmol of Br512 enzyme variants were challenged on a PCR machine that was pre-heated to the temperatures indicated in the figures. After the heat challenges, the tubes were immediately removed from the PCR machine and cooled on an ice-cooled metal rack at least for 5mins. LAMP assay was performed at 65°C for two hours unless otherwise indicated. Fluorescence signals were recorded every 4min in the FAM channel provided by LightCycler 96 software preset.

### Dye-based protein thermal shift assay

The T_m_ (Transition midpoint; Melting Temperature) of the various enzyme variants were measured using Protein Thermal ShiftTM reagents (Thermo Fisher; Catalog Number: 4461146) according to the manufacturer’s instruction. Briefly, a total 40μg (5μg/μL) of each enzyme variants in final dialysis buffer (see Br512 purification protocol) were added into a reaction mixture (20μL) containing 1X Protein Thermal Shift^™^ Buffer and 1X Protein Thermal Shift^™^ Dye. Fluorescence signals were measured in Texas Red channel provided by LightCycler 96 software preset. The red fluorescence change was measured from 37°C to 95°C with 0.1°C/sec ramp speed. The measured values (delta Fluorescence/delta Temperature) were plotted on the graph with a T_m_ calling tool provided by LightCycler 96 analytical software (Roche)

## RESULTS

### Engineering fusion variants of Bst DNAP

We centered our design around the large fragment of DNA polymerase I (Pol I) from *G. stearothermophilus* (*bst*, GenBank L42111.1), which is frequently used for isothermal amplification reactions (9–11). This fragment (hereafter Bst-LF) lacks a 310 amino acid N-terminal domain that is responsible for 5’ to 3’ exonuclease activity, leading to an increased efficiency of dNTP polymerization (12).

Many larger proteins suffer from slow or inefficient folding (13), and removal of the N-terminal portion may lead to increased protein degradation rates, decreased folding speed, and greater instability (14). Indeed, Bst-LF showed low yields upon purification. Therefore, in place of the exonuclease domain we sought to stabilize *Bst* DNAP via a fusion partner, the small F-actin binding protein villin, also referred to as the villin headpiece (5). We further extended HP35 by twelve amino acids to generate (HP47), where the additional amino acids serve to complete an additional alpha helix (N-3) in the structure, which further packs and stabilizes the hydrophobic core. The HP47 tag was added to the amino terminus of the large fragment of Bst-LF, leading to the enzyme we denote as Br512 (**Figure 1a**). Br512 also contains a N-terminal 8x His-tag for immobilized metal affinity chromatography (IMAC; Ni-NTA). The hypothesis that the villin headpiece could serve as an anchor to improve the folding and / or solubility of Bst-LF proved true, and purification of Br512 proved to be much better than for Bst-LF, ultimately yielding 35 mg of homogenous protein per liter (see also purification protocol, below).

**Figure 1.**
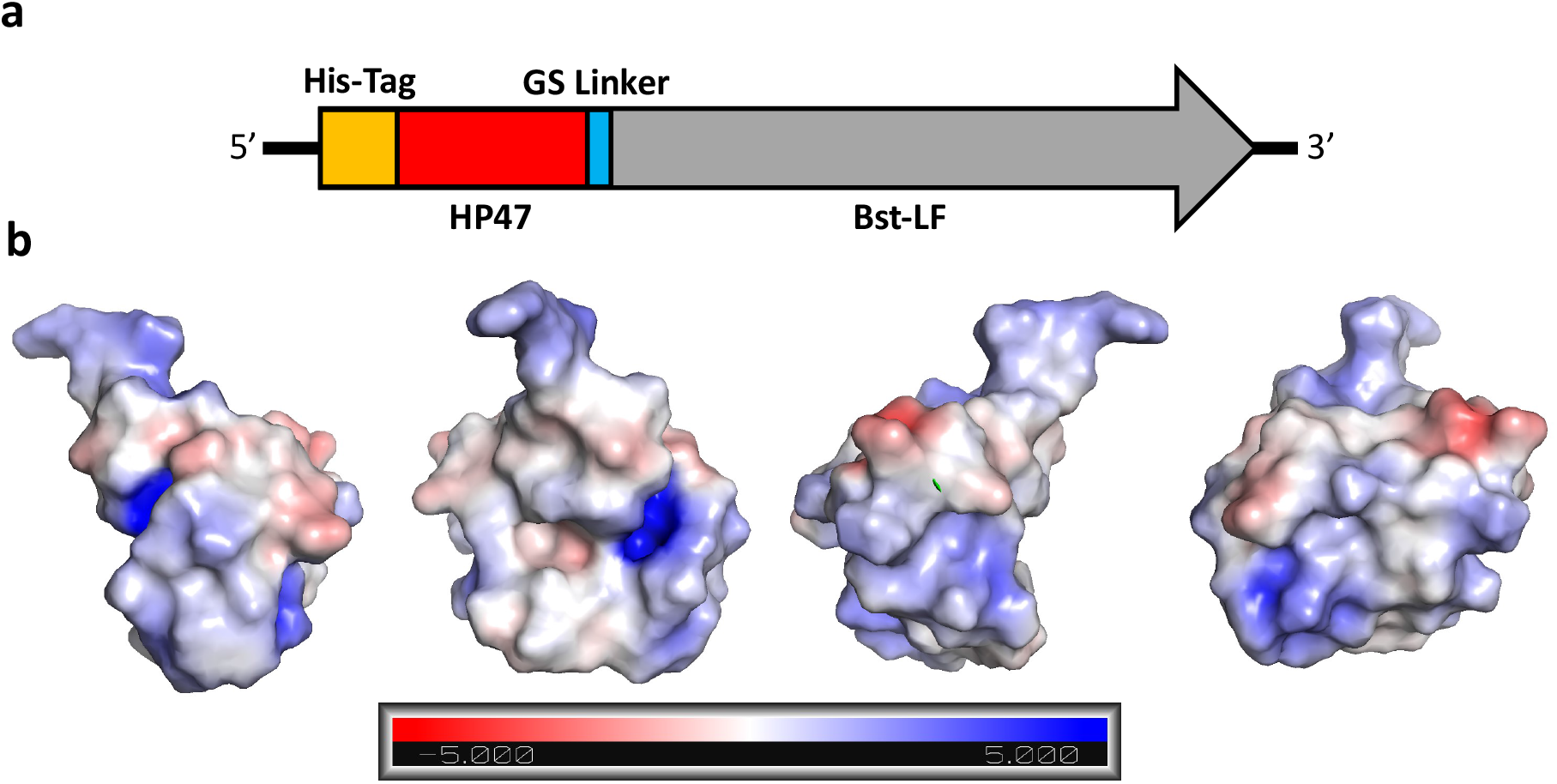
Graphical representation of Br512 and the electrostatic force map of HP47. (a) Br512 was constructed by fusing HP47 with a GS linker to the N-terminal of Bst-LF. A His-Tag was added at the N-terminal of the new fusion protein to aid with purification. (b) Models of HP47 electrostatic force using an Adaptive Poisson-Boltzmann Solver to identify surface charge. The charge designations are referenced in the bar at the bottom. Each graphic is the same model with different orientations rotated on the Y-Axis. Graphics were created in PyMol.

A cluster of positively charged amino acids are known to be crucial for the actin-binding activity of the headpiece domain (15), and thus another potential advantage of the use of HP47 is that it may allow better interactions with nucleic acid templates (7) (**Figure 1b**). In this regard, the HP47 fusion may act similarly to the DNA binding domains used in the construction of synthetic thermostable DNA polymerases that are commonly used for PCR. In the case of Phusion DNA polymerase, the addition of the Sso7d gene, a DNA binding protein from *Sulfolobus solfactaricus*, stabilizes the polymerase/DNA complex and enhances the processivity by up to 9 times, allowing longer amplicons in less time with less influence of PCR inhibitors (4). Similarly, fusion of the *Bst*-like polymerase Gss-polymerase, DNA polymerase I from *Geobacillus* sp. 777, with DNA binding domains from DNA ligase of *Pyrococcus abyssi* or Sto7d protein from *Sulfolobus tokodaii* yielded 3-fold increase in processivity and a 4-fold increase in DNA yield during whole genome amplification (16). We further explored whether a similar improvement might prove true for isothermal amplification by Bst DNA polymerase.

### Performance of Br512 in isothermal amplification assays

The development of isothermal amplification assays that are both sensitive and robust to sampling is key for continuing to mitigate the ongoing coronavirus pandemic (17). To this end, loop-mediated isothermal amplification has proven to be a useful assay for the detection of SARS-CoV-2 (18–21), including in clinical settings (22). However, LAMP is well-known to frequently produce spurious amplicons, even in the absence of template, and thus colorimetric and other methods that do not use sequence-specific probes may be at risk for generating false positive results (23), and we therefore developed oligonucleotide strand displacement probes, that are only triggered in the presence of specific amplicons. These probes are essentially the equivalent of TaqMan probes for qPCR, and can work either in an end-point or continuous fashion with LAMP (23). Base-pairing to the toehold region is extremely sensitive to mismatches, ensuring specificity, and the programmability of both primers and probes makes possible rapid adaptation to the evolution of new SARS-CoV-2 or other disease variants. We have also shown that higher order molecular information processing is also possible, such as integration of signals from multiple amplicons (24).

Our variant of LAMP, which we term LAMP-OSD (for Oligonucleotide Strand Displacement), is designed to be easy to use and interpret, and we have previously shown that it can sensitively and reliably detect SARS-CoV-2, including following direct dilution from saliva (24). Although we have largely mitigated non-specific signaling of LAMP and made it more robust for point of need application, the limited choice and supply, and concomitant expense of LAMP enzymes, constitutes a significant roadblock to widespread application of rapid LAMP-based diagnostics. Br512 presents a potential generally available solution to these issues.

To assess whether the engineered changes introduced in Br512 had an impact on enzyme activity, we first compared the strand displacing DNA polymerase activity of Br512 with that of the wild type Bst-LF enzyme and a commercially-sourced engineered Bst-LF with improved amplification speed, Bst 2.0 (New England Biolabs). We set up duplicate LAMP-OSD assays (23) for the human glyceraldehyde 3-phosphate dehydrogenase (*GAPDH*) gene using either 16 units of Bst 2.0 (typical amount used in most of our LAMP-OSD assays), 20 picomoles (pm) of Bst-LF (a previously optimized amount), or 0.2 pm, 2 pm, 20 pm, or 200 pm of Br512. Real-time measurement of OSD probe fluorescence revealed that in the presence of 6000 template DNA copies the DNA polymerase activity of 20 pm of Br512 was comparable to that of 16 units of Bst 2.0 (**Supplementary Figure 1**). The addition of more Br512 did not yield further improvements, although lower amounts reduced the amplification efficiency. In the absence of specific templates, none of the enzymes generated false OSD signals.

When we set up assays with optimized enzyme amounts and either an optimized buffer system, G6 (developed for reverse transcription reactions with 4 mM (B) or 8 mM (D) Mg2^+^) (25), or Isothermal buffer (provided by New England Biolabs), we found that 20 pm of Br512 per LAMP-OSD assay performed comparably to 16 units of Bst 2.0 in terms of both speed and limit of detection (**Figure 2, panels A, C, and D**). Bst-LF also demonstrated a similar detection limit but with a slower time to signal (**Figure 2, panel B**). Similar results were observed via real-time measurements of amplification kinetics using the fluorescent intercalating dye, EvaGreen in place of sequence-specific OSD probes (**Supplementary Figure 2**). Taken together, these results demonstrate that presence of the villin H47 fusion domain in Br512 improves its speed of amplification relative to Bst-LF bringing it at par with the DNA amplification speed of Bst 2.0.

**Figure 2.**
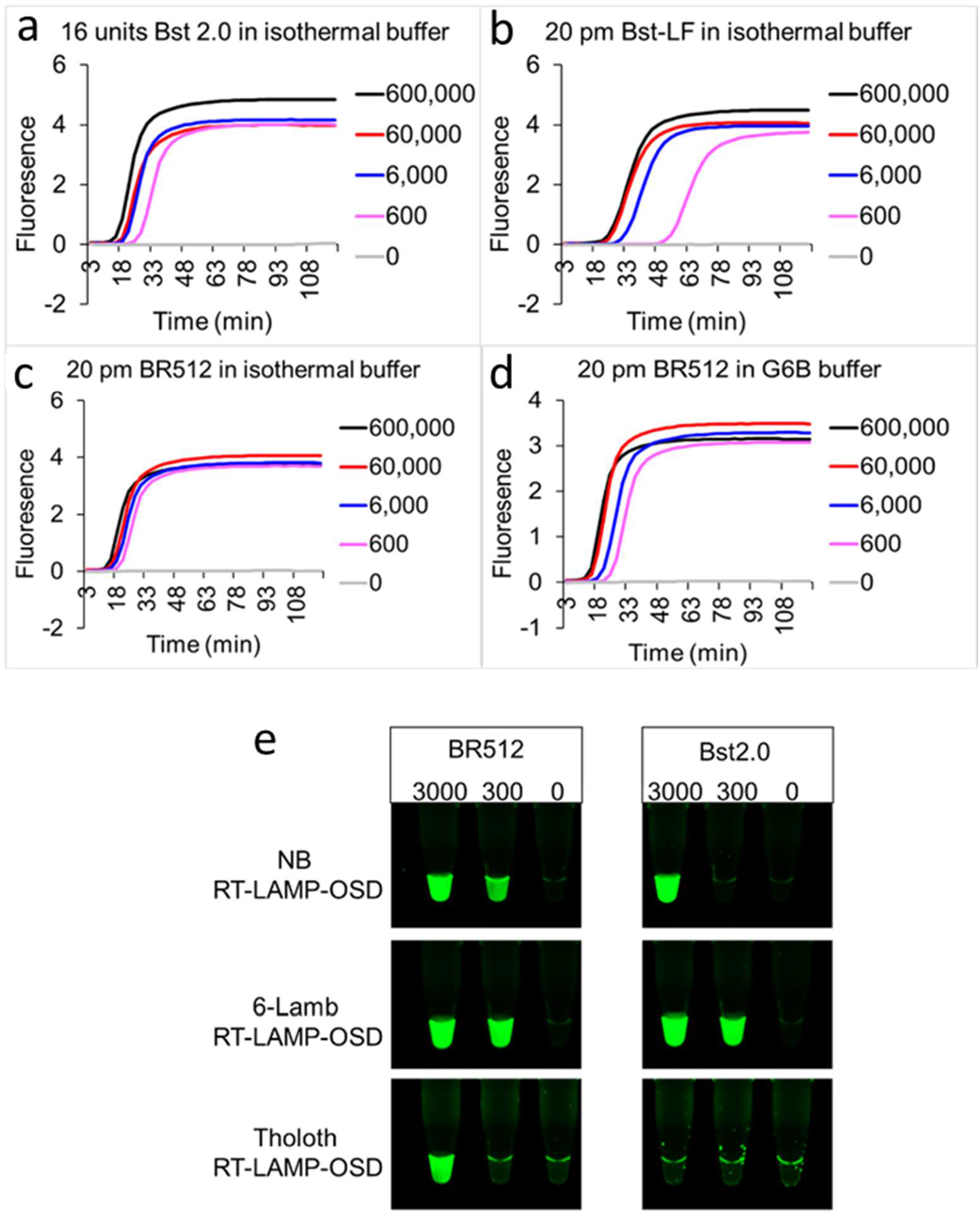
Comparison of Br512, Bst-LF, and Bst 2.0 in LAMP-OSD assays of DNA and RNA templates. LAMP-OSD assays for the human *GAPDH* gene were carried out with 16 units of commercially sourced Bst 2.0 (panel A), 20 pm of in-house purified Bst-LF (panel B), or 20 pm of Br512 (panels C and D) in the indicated reaction buffers. Amplification curves were observed in real-time at 65 °C by measuring OSD fluorescence in reactions seeded with 600,000 (black traces), 60,000 (red traces), 6,000 (blue traces), 600 (pink traces), and 0 (gray traces) copies of *GAPDH* plasmid templates. Three SARS-CoV-2-specific RT-LAMP-OSD assays, NB, 6-Lamb, and Tholoth, that target three different regions in the viral genomic RNA were operated using 20 pm of Br512 (left panels) in 1X G6D buffer or 16 units of commercially sourced Bst 2.0 (right panels) in 1X Isothermal buffer (panel E). SARS-CoV-2 viral genomic RNA templates per reaction are indicated above each column of tubes. Images of OSD fluorescence taken at assay endpoint (after 60 min of amplification at 65 °C followed by cooling to room temperature) are depicted.

Bst DNA polymerase has been described to possess an inherent reverse transcriptase (RT) activity with chemically diverse nucleic acid templates, including ribonucleic acid (RNA), α-l-threofuranosyl nucleic acid (TNA), and 2′-deoxy-2′-fluoro-β-d-arabino nucleic acid (FANA) (FANA) (26,27). Therefore, we tested the performance of Br512 in RT-LAMP-OSD assays in order to determine whether the engineered enzyme could be used for direct amplification from SARS-CoV-2 RNA. We set up duplicate reactions with three different primer sets that we had previously shown worked well with RT-LAMP-OSD (termed NB, Tholoth, and 6-Lamb (24)). These assays were seeded with 3000, 300, or 0 copies of the viral genomic RNA and endpoint OSD fluorescence was imaged following 60 min of amplification at 65 °C. All three RT-LAMP-OSD assays performed using Br512 developed bright green OSD fluorescence in the presence of viral genomic RNA indicating successful reverse transcription and LAMP amplification (**Figure 2E**). While NB and 6-Lamb assays executed with Br512 could readily detect a few hundred viral genomes, the Tholoth assay produced visible signal only with a few thousand viral RNA copies. In contrast, Bst 2.0 demonstrated less robust reverse transcription ability, and failed to generate any OSD signal in the Tholoth assays, both in its companion Isothermal buffer (**Figure 2E)** as well as in the G6D buffer (**Supplementary Figure 3**). Bst 2.0 could reverse transcribe and amplify NB and 6-Lamb sequences resulting in visible OSD fluorescence (**Figure 2E**), but its detection limit for both assays was higher than with Br512. In all cases, in the absence of primers reactions remained dark.

The reliability of detection is an issue, especially at the limit of detection. Br512 could detect 300 SARS-CoV-2 genomes in 80% of NB and 100% of 6-Lamb assays, while Bst 2.0 was successful at detecting this copy number in 25% and 75% of the assays, respectively (**Supplementary Figure 4**). In addition, Br512 generally demonstrated faster amplification kinetics compared to Bst 2.0 (**Supplementary Figure 5**). The superiority of Br512 as an enzyme for RT-LAMP was even more significant when compared to Bst-LF. The wild-type, parental enzyme failed to amplify the Tholoth RNA sequence, while only detecting SARS-CoV-2 genomic RNA in 16% of NB assays and 33% of 6-Lamb assays (**Supplementary Figure 4**). Bst-LF also demonstrated slower amplification kinetics compared to both Br512 and Bst 2.0 (**Supplementary Figure 5**).

### Machine learning predictions improve Br512 function

Torng and Altman (2017) developed a novel 3D convolutional neural net to examine whether the microenvironments of individual amino acids could be used to classify what amino acids were present throughout a protein’s structure. This algorithm trained across the PDB, and in the end was able to re-predict the identity of a given amino acid in a protein with ca. 40% accuracy. We have since improved upon this algorithm by including additional filters for features such as the presence of hydrogen atoms, partial charge, and solvent accessibility, and have ultimately improved the accuracy of re-prediction to upwards of 70% (28). We were interested in the approximately 30% of amino acids that were not predicted to be wild-type; while this might have merely reflected the inaccuracy of the neural network, it was also possible that nature itself was ‘underpredicting’ the fit of a given amino acid to its microenvironment, and that the ensemble of predicted non-wild-type amino acids at a given position might represent opportunities for mutation. To this end, we instantiated the ability to predict underperforming wild-type amino acid residues in proteins by predicting either positions or precise mutations that led to improvements in function across a variety of proteins, including blue fluorescent protein and phosphomannose isomerase (28). It should be noted that such predictions were not for a particular functionality, such as stability or catalysis, but merely for goodness of fit, with improvements in stability and catalysis being empirical outcomes of those predictions.

With this track record of success, we attempted to apply the improved algorithm, dubbed MutCompute, to predicting amino acids in the *Bst* LF that might benefit from mutation. Initially, residues that interacted directly with DNA were filtered from consideration, as the algorithm has not yet been optimized for protein:nucleic acid interactions. An amino acid-by-amino acid assessment of the remaining residues in the protein was carried out, and those positions that revealed the wild-type residue to be the least fit for a given position were called, and ten substitutions were then made that represented the predicted most fit amino acid (**Figure 3a**).

**Figure 3.**
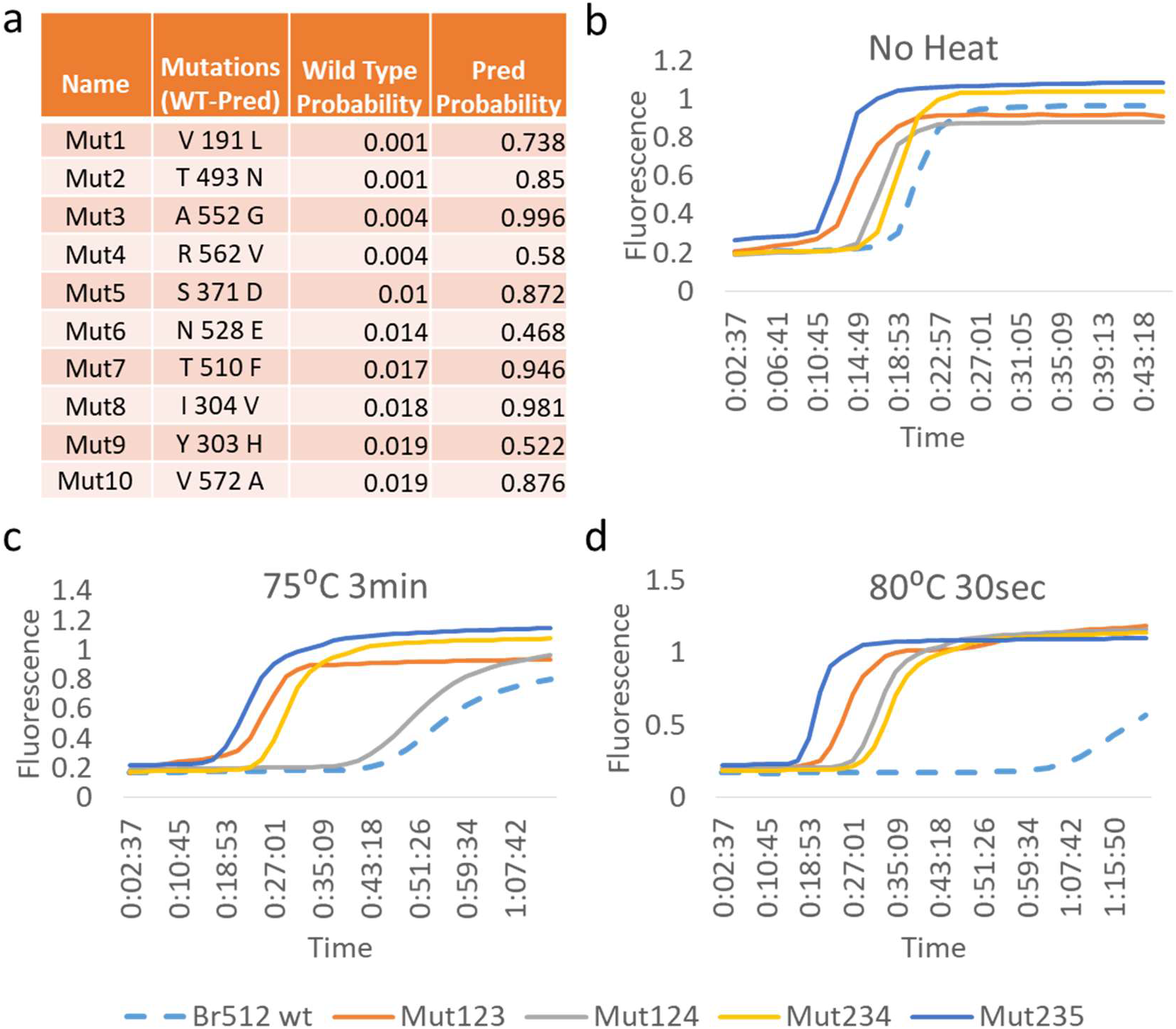
Br512 MutCompute mutations and their effect on enzyme thermal stability. (a) Table listing Br512 stabilizing amino acid substitutions suggested by MutCompute. Wild type (WT) vs predicted (Pred) amino acid mutations (column 2; positions in reference to PDB database ID; 3TAN) were designated as Mut1 to Mut10 according to their predicted priorities. The calculated probabilities of the wild type and predicted amino acids at each position are indicated in columns 3 and 4, respectively. (b-d) Effect of thermal challenge on wildtype and triple mutant MutCompute variants of Br512. Identical *GAPDH* LAMP assays assembled using the same amount of indicated enzymes were subjected to either no heat challenge (b), 3 min at 75°C (c), or 30 sec at 80°C (d) prior to real time measurement of *GAPDH* DNA amplification kinetics at 65°C. Representative amplification curves generated by measuring increases in EvaGreen dye fluorescence (Y-axis) over time (X-axis; time in hh:mm:ss) are depicted as dotted blue (Br512 wild type), burnt orange (Mut123), gray (Mut124), yellow (Mut234), and blue (Mut235) traces.

Surprisingly, of the ten amino acid substitutions suggested by MutCompute, only two (Mut6 and Mut9) showed little or no activity in a standard LAMP assay targeting the gene for human *GAPDH*, while the top 5 (Mut1-5) showed activities as good as or better than the parent enzyme (**Supplementary Figure 7**).

Because we were hoping to introduce additive substitutions and achieve higher thermostability, we also adapted LAMP to serve a simple screen for improved activity at higher temperatures. Initially, enzymes were challenged at temperatures above those typically used for LAMP (75°C and 80°C), before carrying out LAMP reactions at their normal temperature (65°C). Mut1-5 were further assayed with a heat challenge, to determine if they had imparted additional stability to the polymerase (**Supplementary Figure 8**), and both Mut2 and Mut 3 were found to be more thermotolerant than the parental enzyme.

In keeping with our initial hypothesis that different engineering tacks would yield substitutions that had additive impacts on phenotypes, we have previously had great success in combining individual mutations predicted via machine-learning approaches to generate proteins with much higher activities; for example, multiple slightly improved variants of blue fluorescent protein could be combined to yield a variant with 5-fold greater fluorescence, while multiple slightly improved variants of a phosphomannose isomerase could be combined to yield a variant with 5-fold greater solubility (28). Therefore, we examined combinations of the point mutations that showed the greatest activity. We initially generated all possible double combinations of the Muts 1-4 (Mut12, 13, 14, 23, 24, and 34) and carried out LAMP assays and thermal challenges (**Supplementary Figure 9**). Mut23 yielded the most robust activity, in keeping with the results of the initial thermal challenges.

We finally generated four additional triple mutations (Mut123, Mut124, Mut234, Mut235) centered on Mut23 (**Figure 3**). All four triple mutations examined showed robust performance in the normal *GAPDH* LAMP assay (**Figure 3b, Supplementary Figure 10**), and the combined machine-learning predicted mutations also displayed strong thermotolerance relative to the parental enzyme, which itself was already superior to Bst-LF (**Figure 3c, 3d, Supplementary Figure 10**). Mut235 showed the highest activity, and was therefore used as a platform for further, additive engineering. Interestingly, Mut5 on its own has an inactive phenotype at higher temperatures, and seems to serve as a potentiating mutation for additional substitutions.

### Supercharging of the villin headpiece (vHP47) improves Br512 function

Given that the villin headpiece likely assists with contacting nucleic acid substrates via positively charged patches, we sought to further improve this feature via the addition of excess positively charged amino acids (supercharging; (29)). We also anticipated that supercharging would improve the folding, solubility, and stability of the protein, by decreasing the propensity to aggregate, as we and others have previously demonstrated (30–32). We further anticipated that improvements gained via supercharging would be additive with substitutions introduced via machine-learning approaches, as supercharging targets additional biophysical mechanisms for stabilization, such as potentially improving interactions with the DNA substrate.

Since vHP47 is naturally a ‘zwitterionic’ or ‘Janus’ protein that contains two oppositely charged surfaces (**Figure 4a**), we hypothesized that it could be further engineered to orient the domain for binding to nucleic acids and thereby potentially enhancing polymerase activity, as previous nucleic acid binding domains (e.g., Phusion) had done. To this end, we initially surveyed highly solvent-exposed amino acids in the vHP47 crystal structure (PDB: 1YU5), and assessed the relative conservation amongst the top vHP47 100 orthologues from the search (**Supplementary Figure 10**). Variable amino acid positions were selected as candidates for mutagenesis (**Figure 4a, Supplementary Figure 11**), and we also sought to focus supercharging on single surfaces of vHP47 (**Figure 4b**). Eight point mutations (SC1-8; **Figure 4c**) were initially engineered. Four of the amino acid substitutions (SC1-4) were designed to enhance the negatively charged surface, while the other four substitutions (SC5-8) should enhance the opposing, positively charged surface (**Figure 4b**)

**Figure 4.**
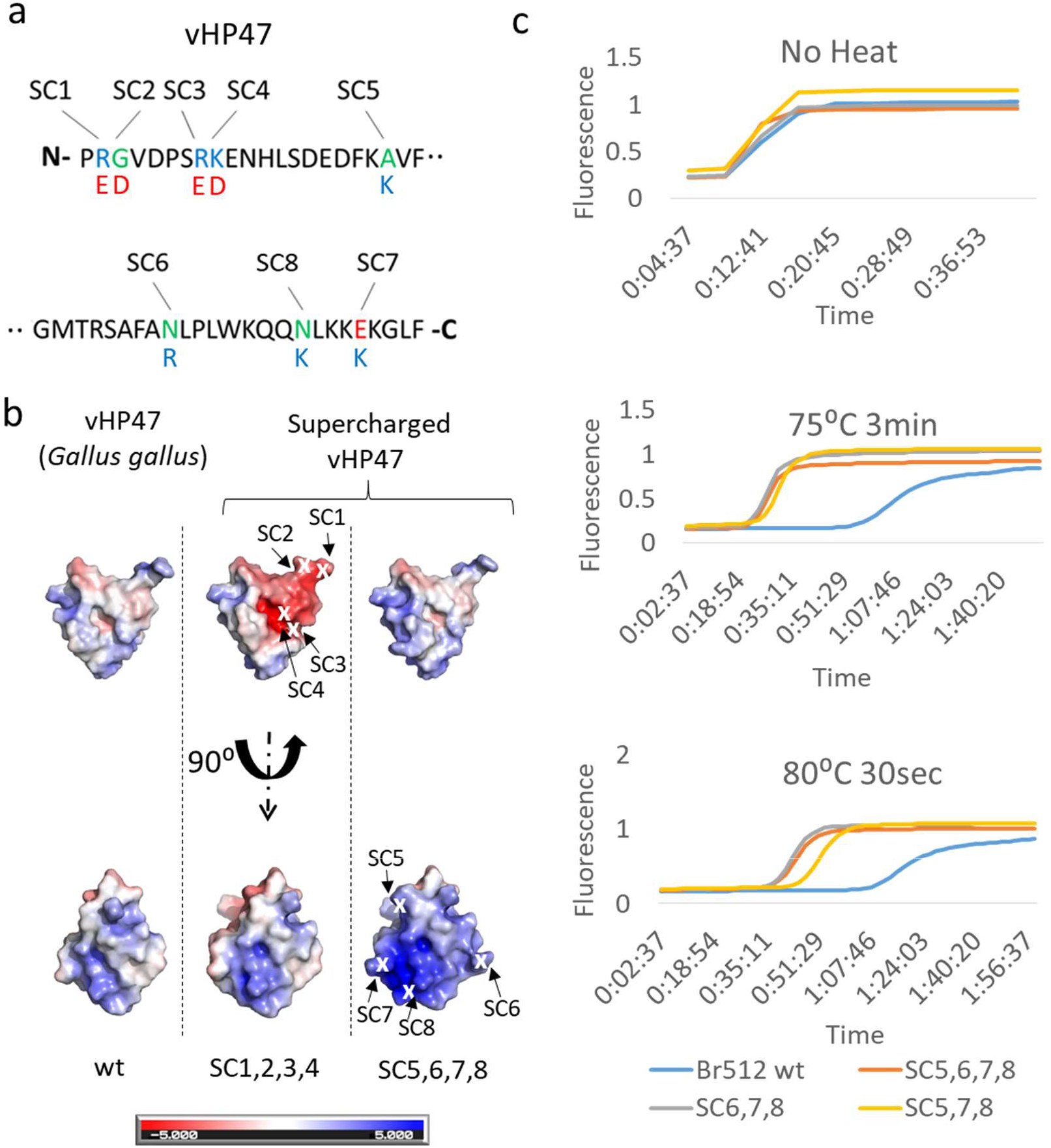
Effect of supercharged villin headpiece on Br512 thermostability. (a) The Villin headpiece (vHP47) amino acid sequence and its corresponding supercharging mutations. Neutral, negatively charged, and positively charged amino acids are depicted by green, red, and blue letter designations, respectively. (b) Surface charge models of wildtype (wt) vHP47 domain and its supercharged variants generated as described in Figure 1. A total eight amino acids of vHP47 designated as SC1-8 were mutated into either negatively (SC1,2,3,4) (Aspartate D/Glutamate E) or positively charged amino acids (SC5,6,7,8) (Lysine; K/Arginine; R). (c) Effect of thermal challenge on triple and quadruple positively supercharged mutants of Br512. Identical *GAPDH* LAMP assays assembled using the same amount of indicated enzymes were subjected to either no heat challenge (top panel), 3 min at 75°C (middle panel), or 30 sec at 80°C (bottom panel) prior to real time measurement of *GAPDH* DNA amplification kinetics at 65°C. Representative amplification curves generated by measuring increases in EvaGreen dye fluorescence (Y-axis) over time (X-axis; time in hh:mm:ss) are depicted as blue (Br512 wild type), burnt orange (SC5,6,7,8), gray (SC6,7,8), and yellow (SC5,7,8) traces. The effect of various single, double, and triple mutations are shown in Supplementary Figures 12 and 13.

While the four, individual negatively charged substitutions exhibited no enhanced activity, the four positively charged substitutions (SC5, SC6, SC7, SC8) showed enhanced activities under heat challenges (**Supplementary Figure 12**). To examine the combined effect of the four mutations, we generated various double and triple combinations (**Supplementary Figure 13**). The higher order mutations showed additive activities with heat challenges, suggesting that the increased surface charge imparts either greater overall activity or stability to vHP47 (**Figure 4c**; **Supplementary Figure 13, 14**). The two best variants (SC5,6,7,8 and SC6,7,8) showed exponential target amplification starting as early as 18 mins (after a heat challenge at 75°C) and 33 mins (after a heat challenge at 80°C), both of which were at least a half an hour earlier than was seen with the wild-type enzyme at similar temperatures (**Figure 4c, Supplementary Figure 14**).

### Combining mutations further improves enzyme function

As the two independent efforts to engineer the Br512 enzyme resulted in a great success, we sought to examine combination of the mutations on the Br512. In this regard, we were also tacitly testing the hypothesis that attempts to promote different aspects of enzyme stability (fusion domains, improved amino acid interactions, and supercharging to promote folding and DNA interactions) would operate independently and additively. This hypothesis seemed especially reasonable given that mutations were introduced into separate portions of the Br512 enzyme (Bst-LF and vHP47, respectively).

Using the engineered Mut235 variant as a platform, we added the best, combined supercharged mutations (SC678 and SC5678) to generate Mut235-SC678 (renamed as Br512g3.1) and Mut235-SC5678 (renamed as Br512g3.2; **Figure 5**). LAMP assays were performed with these two combined variants, and their performance was quantitated by looking at amplification threshold cycle (C_t_) values, which indicate the time to a significant signal above background. These two variants had C_t_ values of 10.8 mins, which is approximately 5 mins faster than the Br512 wt (Ct of 15.6 mins) even in the absence of a heat challenge prior to LAMP assay (**Figure 5a**). As anticipated, the combined variants also showed the most robust target amplification after heat challenges (**Figure 5b-d**). For instance, the combined variants were faster (10.1mins) to yield a signal than either of the parental mutations (13 mins and 16.2 mins for Mut235 and SC678, respectively), and at least 24 minutes faster than the parental Br512 enzyme (**Figure 5b**). After an 80°C heat challenge the combined variants showed further improvements relative to either the parental mutations (Mut235, SC678, SC5678) or to the parental enzyme (**Figure 5c**). Overall, the combined variants Br512g3.1 and Br512g3.2 remained largely active up to 82°C (**Figure 5d**), a temperature that completely inactivated the parental enzyme.

**Figure 5.**
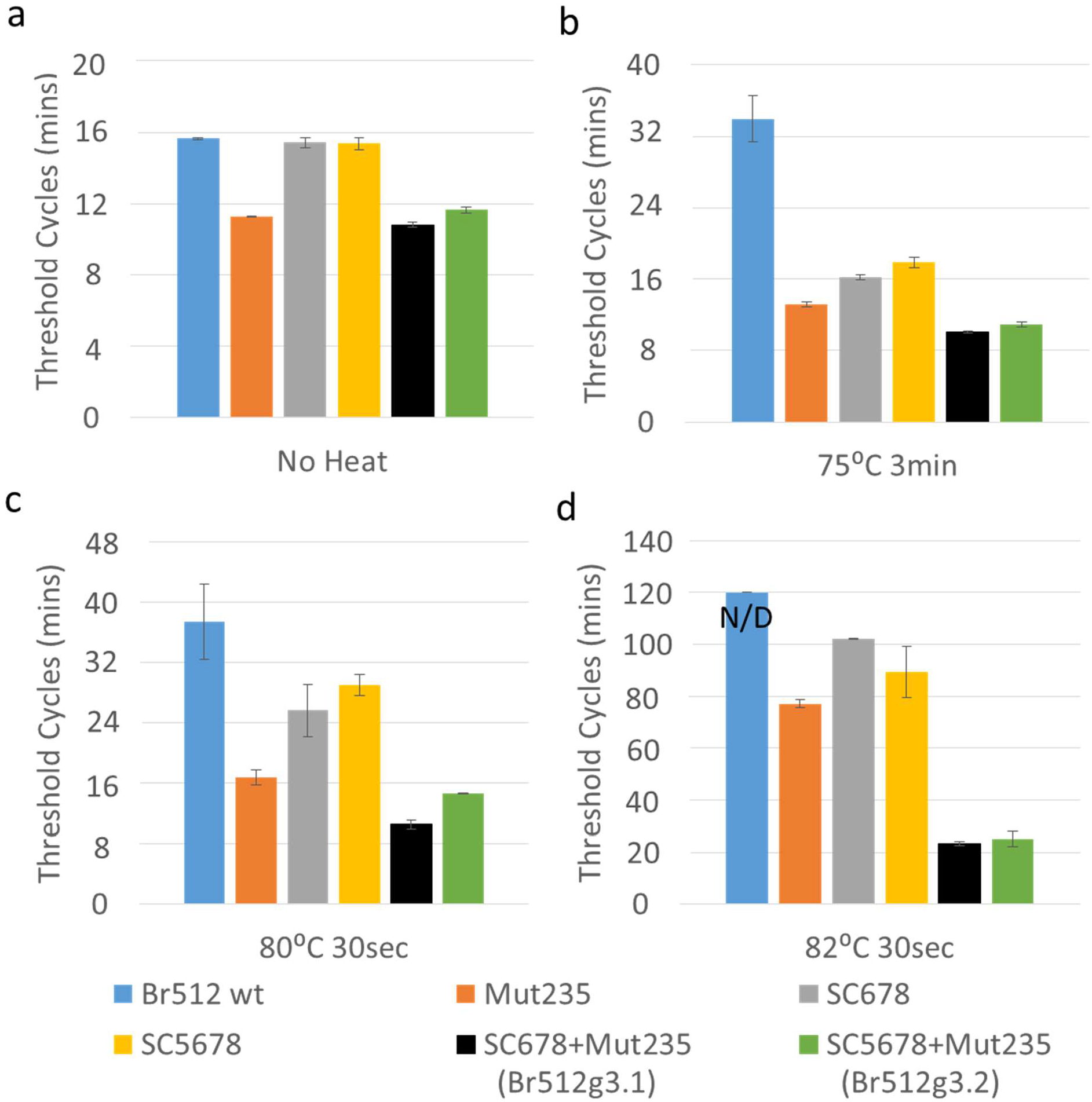
Effect of combining MutCompute and supercharging mutations on Br512 thermal stabilities. Identical *GAPDH* LAMP assays assembled using either wildtype (wt), Supercharged-villin headpiece (SC), MutCompute (Mut), or combined SC+Mut Br512 variants were subjected to either no heat challenge (a), 3 min at 75 °C (b), 30 sec at 80 °C (c), or 30 sec at 82 °C (d) prior to real time measurement of *GAPDH* DNA amplification kinetics at 65°C. Threshold cycle (mins; time to detection) values for amplification of 20 pg (6×10^7^ copies) *GAPDH* DNA templates were calculated using the LightCycler 96 software and plotted as bar graphs (Blue: Br512 wild type; Burnt orange: Mut235; Gray: SC678; Black: SC678+Mut235; Green: SC5678+Mut235). Time to reach the threshold cycles (mins; time to detection) are shown as bar graphs, in minutes. Standard deviation in C_t_ values calculated from three replicate experiments is depicted as error bars. N/D= Amplification not detected.

While the increased thermostability of the engineered variants was strongly indicated by the thermal challenge assays, we also used a dye-based protein thermal shift assay (TSA) to determine the melting temperatures of the proteins (**Supplementary Figure 15**). The parental enzyme Br512 showed a slightly higher T_m_ value (78.4°C) compared to its parental enzyme Bst-LF (78.2°C), but the two best variants showed greatly improved T_m_ values: 80.4 C and 80.6°C for SC5678-Mut235, (renamed as Br512g3.1) and SC678-Mut235 (renamed as Br512g3.2), respectively, further supporting the enhanced thermostability of the engineered variants.

### Combined variants allow extraordinarily fast LAMP reactions

The speed of LAMP reactions is generally limited by the strand displacement ability of the enzyme, and by the related ability of the amplified DNA to form single-strands that can fold back and create a new 3’ hairpin in the growing concatemers. Given the increases in speed observed during thermal challenges, it seemed possible that we could identify an optimal temperature for ultrafast amplification, where the strand displacement capabilities of the combined variants would be optimally balanced with the ability to bind primers and form new 3’ hairpins. By scanning through several temperatures (**Supplementary Figure 16**), we found that the LAMP reaction could enter into a readily detected exponential phase as soon as 6 minutes, with the overall reaction being completed by ~10min at 74°C LAMP (**Figure 6a**). Quantitation by time to detection (threshold cycle) showed that the C_t_ value of the variants was as low as ~6mins (**Figure 6b**). Overall, the time to detection (reaction time to reach C_t_) for 6×10^7^ copies of *GAPDH* DNA templates was ~6 min (**Figure 6a**), ~7 min for Br512g3.1 and g3.2, respectively for 600,000 copies (**Figure 6c**), and ~9 min for 6,000 copies (**Figure 6c**). In contrast, the parental Br512 enzyme was completely inactivated in continuous amplification assays (as opposed to thermal challenges) at 73°C and 74°C (**Supplementary Figure 16, and Figure 6a,b,c**). The specificities of the *GAPDH* LAMP assays shown in this study were verified with non-template controls (**Supplementary Figure 17 and 18**). In general, the engineered enzymes are more thermotolerant to brief challenges than to continuous function at high temperatures; for example, enzymes that are thermotolerant for short periods of time at 75°C or 80°C may not show activity in LAMP at 74°C.

**Figure 6.**
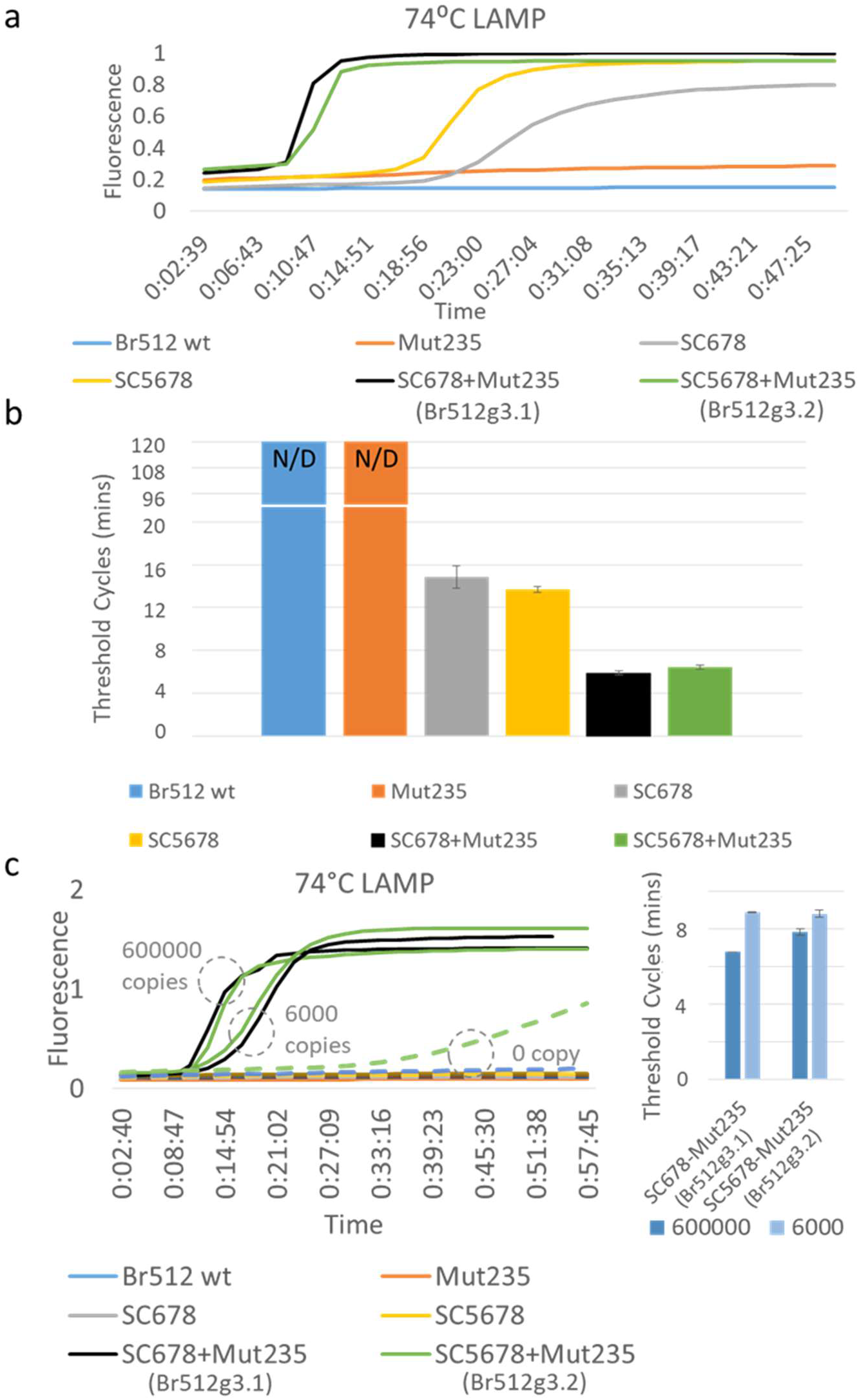
Comparison of Br512 variants in high temperature LAMP assays. (a-b) Identical *GAPDH* LAMP assays were assembled using the same amounts of either wildtype (blue) or mutant Br512 variants (burnt orange: Mut235; gray: SC678; yellow: SC5678; black: SC678+Mut235 (Br512g3.1); green: SC5678+Mut235 (Br512g3.2)) and incubated at 74°C for up to two hours. Amplification kinetics of *GAPDH* DNA templates were determined by real time measurement of EvaGreen dye fluorescence and the threshold cycles (mins; time to detection) for amplification of 20 pg (6×10^7^ copies) *GAPDH* DNA templates were calculated using the Lightcyler 96 software. (c) (Left panel) High temperature (74°C) LAMP assay amplification results from 600,000; 6,000; and 0 copies of *GAPDH* DNA templates are shown. Dotted lines indicate traces from 0 templates (Right panel) Calculated threshold cycles (mins; time to detection) from LAMP assays shown in (c). Representative changes in fluorescence intensities (Y-axis) over time are depicted as amplification curves (Panel a and c) while average C_t_ values from three replicate experiments are plotted as bar graphs (Panel b: N/D= Amplification Not Detected, Error Bar=S.D. n=3).

Finally, we performed a 73°C LAMP assay using commercially available Bst2.0 and Bst3.0 enzymes (from NEB) with the buffer (Isothermal II buffer) provided by the manufacturer. The top two variants outperformed Bst2.0 and Bst3.0 polymerases in the *GAPDH* LAMP assay (**Supplementary Figure 19)**, even in the alternative buffer. While Bst2.0 was completely inactive, Bst3.0 was slower to produce a signal than our top two variants (Br512g3.1 and Br512g3.2) by about 2 mins.

## DISCUSSION

There have been various approaches advanced for engineering DNA polymerases for improved function in a variety of settings (36,37), but these typically focus on a single method for improvement or a single property of the polymerase. It has been hypothesized that impacting different kinetic properties of a protein can lead to additive improvements in protein function (1,2), a hypothesis that also implicitly underlies the many highly successful examples in which DNA shuffling is used to improve protein function (38,39). We and others have previously shown that thermal stability is a global property of a protein, and that amino acid changes throughout a protein’s structure can lead to a higher melting temperature and performance at higher temperatures (40,41), and we therefore attempted to improve the function of Bst-LF at increasingly higher temperatures through several different, complementary mechanisms. The addition of the ultra-fast folding domain HP47 was anticipated to assist with formation of protein tertiary structure following translation (so-called ‘assisted folding;’ (42–44)) and improve the solubility of the enzyme for purification (43,45), but what was unanticipated was that HP47 might also allow the polymerase to better interact with DNA via its zwitterionic nature, and thereby improve the ability to carry out LAMP reactions. By further using machine-learning methods and supercharging we sought to further improve the stability and functionality of the enzyme in additive ways (46). In fact, mutations introduced at multiple sites around the enzyme and its fusion partner generally proved additive, as has previously been observed for structurally distant mutations in other proteins, including transcription factors (47), kinesin (48), and serine proteases (49). Ultimately, we were eventually able to generate enzymes (Br512g3.1 and Br512g3.2) that could perform high temperature (74°C), ultra-fast (6 minute) LAMP reactions, and still provide reliable and consistent outputs. A great advantage of high temperature LAMP seems to be reduced background, and as the Br512g3 enzymes make possible exploration of these new high temperature reactions it is likely that primers and probes will also have to be concomitantly redesigned. Overall, these combined enzyme engineering efforts provide a blueprint for how to generally improve molecular biology enzymes, and generate multiple new options for carrying out and translating LAMP reactions into practice, including in point-of-care settings.

## DATA AVAILABILITY

Full sequence and annotations of the pKAR2-Br512 plasmid are available in Supplementary Table 2. The pKAR2-Br512 expression plasmid is readily available for distribution from Addgene. (https://www.addgene.org/161875/)

## FUNDING

This work was supported by grants from National Science Foundation (2027169), National Institutes of Health (1R01EB027202-01A1, 3R01EB027202-01A1S1), The Welch Foundation (F-1654), and National Aeronautics and Space Administration (NNX15AF46G).

## TABLE AND FIGURES LEGENDS

**Figure 1. Graphical representation of Br512 and the electrostatic force map of HP47.** (a) Br512 was constructed by fusing HP47 with a GS linker to the N-terminal of Bst-LF. A His-Tag was added at the N-terminal of the new fusion protein to aid with purification. (b) Models of HP47 electrostatic force using an Adaptive Poisson-Boltzmann Solver to identify surface charge. The charge designations are referenced in the bar at the bottom. Each graphic is the same model with different orientations rotated on the Y-Axis. Graphics were created in PyMol.

**Figure 2. Comparison of Br512, Bst-LF, and Bst 2.0 in LAMP-OSD assays of DNA and RNA templates.** LAMP-OSD assays for the human *GAPDH* gene were carried out with 16 units of commercially sourced Bst 2.0 (panel A), 20 pm of in-house purified Bst-LF (panel B), or 20 pm of Br512 (panels C and D) in the indicated reaction buffers. Amplification curves were observed in real-time at 65 °C by measuring OSD fluorescence in reactions seeded with 600,000 (black traces), 60,000 (red traces), 6,000 (blue traces), 600 (pink traces), and 0 (gray traces) copies of *GAPDH* plasmid templates. Three SARS-CoV-2-specific RT-LAMP-OSD assays, NB, 6-Lamb, and Tholoth, that target three different regions in the viral genomic RNA were operated using 20 pm of Br512 (left panels) in 1X G6D buffer or 16 units of commercially sourced Bst 2.0 (right panels) in 1X Isothermal buffer (panel E). SARS-CoV-2 viral genomic RNA templates per reaction are indicated above each column of tubes. Images of OSD fluorescence taken at assay endpoint (after 60 min of amplification at 65 °C followed by cooling to room temperature) are depicted.

**Figure 3. Br512 MutCompute mutations and their effect on enzyme thermal stability.**

(a) Table listing Br512 stabilizing amino acid substitutions suggested by MutCompute. Wild type (WT) vs predicted (Pred) amino acid mutations (column 2; positions in reference to PDB database ID; 3TAN) were designated as Mut1 to Mut10 according to their predicted priorities. The calculated probabilities of the wild type and predicted amino acids at each position are indicated in columns 3 and 4, respectively. (b-d) Effect of thermal challenge on wildtype and triple mutant MutCompute variants of Br512. Identical *GAPDH* LAMP assays assembled using the same amount of indicated enzymes were subjected to either no heat challenge (b), 3 min at 75°C (c), or 30 sec at 80°C (d) prior to real time measurement of *GAPDH* DNA amplification kinetics at 65°C. Representative amplification curves generated by measuring increases in EvaGreen dye fluorescence (Y-axis) over time (X-axis; time in hh:mm:ss) are depicted as dotted blue (Br512 wild type), burnt orange (Mut123), gray (Mut124), yellow (Mut234), and blue (Mut235) traces.

**Figure 4. Effect of supercharged villin headpiece on Br512 thermostability.**

(a) The Villin headpiece (vHP47) amino acid sequence and its corresponding supercharging mutations. Neutral, negatively charged, and positively charged amino acids are depicted by green, red, and blue letter designations, respectively. (b) Surface charge models of wildtype (wt) vHP47 domain and its supercharged variants generated as described in Figure 1. A total eight amino acids of vHP47 designated as SC1-8 were mutated into either negatively (SC1,2,3,4) (Aspartate D/Glutamate E) or positively charged amino acids (SC5,6,7,8) (Lysine; K/Arginine; R). (c) Effect of thermal challenge on triple and quadruple positively supercharged mutants of Br512. Identical *GAPDH* LAMP assays assembled using the same amount of indicated enzymes were subjected to either no heat challenge (top panel), 3 min at 75°C (middle panel), or 30 sec at 80°C (bottom panel) prior to real time measurement of *GAPDH* DNA amplification kinetics at 65°C. Representative amplification curves generated by measuring increases in EvaGreen dye fluorescence (Y-axis) over time (X-axis; time in hh:mm:ss) are depicted as blue (Br512 wild type), burnt orange (SC5,6,7,8), gray (SC6,7,8), and yellow (SC5,7,8) traces. The effect of various single, double, and triple mutations are shown in **Supplementary Figures 12 and 13**.

**Figure 5. Effect of combining MutCompute and supercharging mutations on Br512 thermal stabilities.** Identical *GAPDH* LAMP assays assembled using either wildtype (wt), Supercharged-villin headpiece (SC), MutCompute (Mut), or combined SC+Mut Br512 variants were subjected to either no heat challenge (a), 3 min at 75 °C (b), 30 sec at 80 °C (c), or 30 sec at 82 °C (d) prior to real time measurement of *GAPDH* DNA amplification kinetics at 65°C. Threshold cycle (C_t_) values for amplification of 20 pg (6×10^7^ copies) *GAPDH* DNA templates were calculated using the LightCycler 96 software and plotted as bar graphs (Blue: Br512 wild type; Burnt orange: Mut235; Gray: SC678; Black: SC678+Mut235; Green: SC5678+Mut235). Time to reach the threshold cycles (C_t_) are shown as bar graphs, in minutes. Standard deviation in C_t_ values calculated from three replicate experiments is depicted as error bars. N/D= Amplification not detected.

**Figure 6. Comparison of Br512 variants in high temperature LAMP assays.** (a-b) Identical *GAPDH* LAMP assays were assembled using the same amounts of either wildtype (blue) or mutant Br512 variants (burnt orange: Mut235; gray: SC678; yellow: SC5678; black: SC678+Mut235 (Br512g3.1); green: SC5678+Mut235 (Br512g3.2)) and incubated at 74°C for up to two hours. Amplification kinetics of *GAPDH* DNA templates were determined by real time measurement of EvaGreen dye fluorescence and the threshold cycles (C_t_) for amplification of 20 pg (6×10^7^ copies) *GAPDH* DNA templates were calculated using the Lightcyler 96 software. (c) (Left panel) High temperature (74°C) LAMP assay amplification results from 600,000; 6,000; and 0 copies of *GAPDH* DNA templates are shown. Dotted lines indicate traces from 0 templates (Right panel) Calculated threshold cycles (C_t_) in minutes from LAMP assays shown in (c). Representative changes in fluorescence intensities (Y-axis) over time are depicted as amplification curves (Panel a and c) while average C_t_ values from three replicate experiments are plotted as bar graphs (Panel b; 1ΔCt = 4mins; N/D= Amplification Not Detected; Error Bar=S.D. n=3).

## Notes

### Competing Interest Statement

The authors have declared no competing interest.

