## Supplementary Figure 1-20, Table1-2 for "Multi-modal engineering of *Bst* DNA polymerase for thermostability in ultra-fast LAMP reactions"

### Supplementary Information

Supplementary Figure 1
Supplementary Figure 2
Supplementary Figure 3
Supplementary Figure 4
Supplementary Figure 5
Supplementary Figure 6
Supplementary Figure 7
Supplementary Figure 8
Supplementary Figure 9
Supplementary Figure 10
Supplementary Figure 12
Supplementary Figure 13
Supplementary Figure 14
Supplementary Figure 15
Supplementary Figure 16
Supplementary Figure 17
Supplementary Figure 18
Supplementary Figure 19
Supplementary Figure 20

Supplementary Table 1
Supplementary Table 2

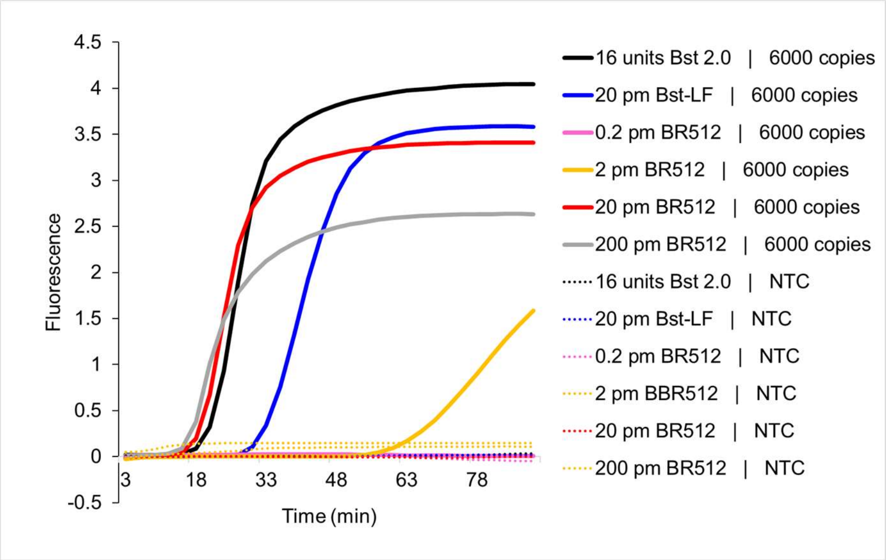


**Supplementary Figure 1. Effect of varying amounts of Br512 on LAMP-OSD of DNA templates.** Indicated amounts of Br512 were compared with indicated amounts of in-house purified Bst-LF and commercially sourced Bst 2.0 in human *GAPDH* gene-specific LAMP-OSD assays operated in 1X isothermal buffer (NEB). Reactions were seeded with either 6000 copies of *GAPDH* plasmid template or with no specific templates (NTC). Amplification curves generated by real-time measurement of OSD fluorescence at 65 °C are depicted.

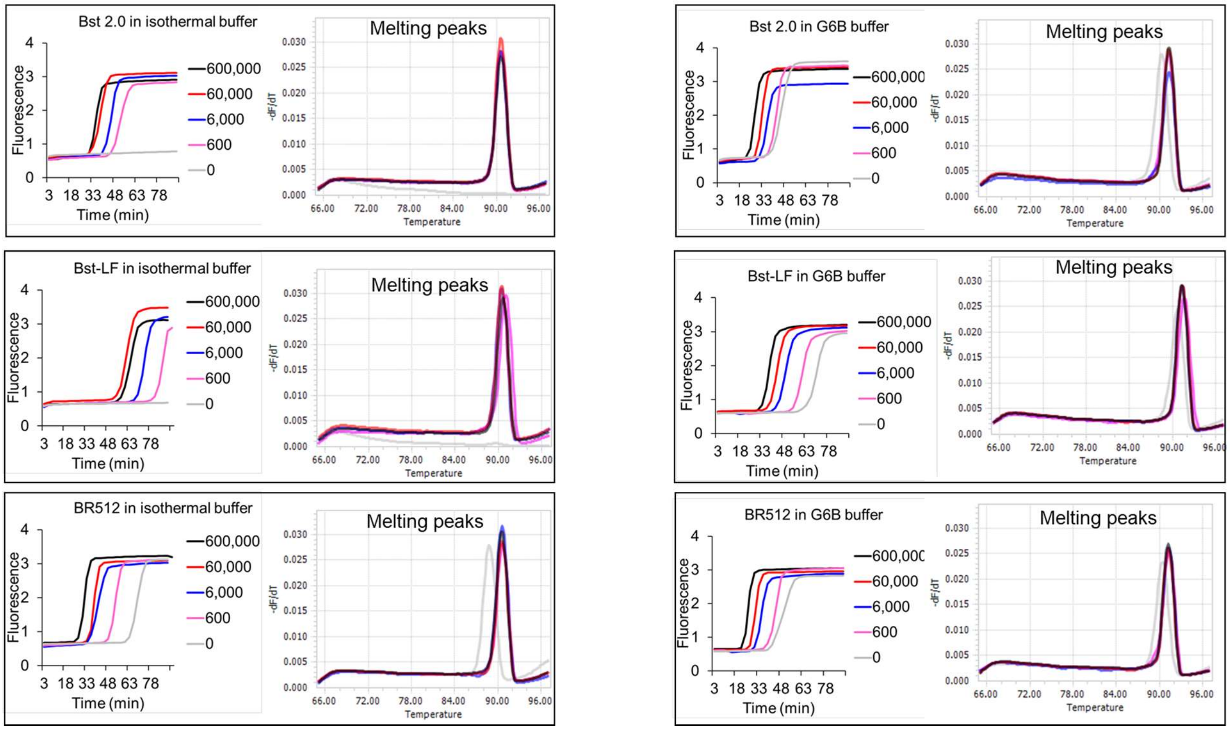


**Supplementary Figure 2. Comparison of Br512, Bst-LF, and Bst 2.0 in LAMP assays of DNA templates read using EvaGreen intercalating dye.** LAMP assays for human *GAPDH* gene were operated using Bst 2.0, Bst-LF, or Br512 in indicated reaction buffers. Amplification curves observed in real-time at 65 °C by measuring EvaGreen fluorescence in reactions seeded with 600,000 (black traces), 60,000 (red traces), 6,000 (blue traces), 600 (pink traces), and 0 (gray traces) copies of *GAPDH* plasmid templates are depicted. LAMP amplicons were analyzed using the 'melt curve analysis' on LightCycler 96 real-time PCR machine and resulting melting peaks are indicated in the corresponding colored traces.

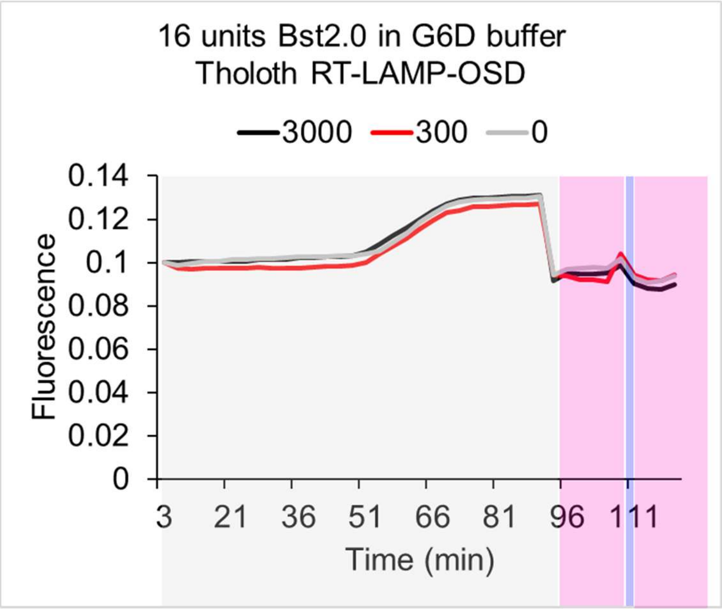


**Supplementary Figure 3. Bst 2.0 Tholoth RT-LAMP-OSD assay executed in G6D buffer.** Tholoth RT-LAMP-OSD assays for SARS-CoV-2 were operated using Bst 2.0 in G6D reaction buffer. OSD fluorescence measured in real-time during assay incubation at 65 °C are depicted within gray shaded boxes for reactions seeded with 3,000 (black traces), 300 (red traces), or 0 (gray traces) copies of SARS-CoV-2 genomic RNA templates. Post-amplification phase OSD signal measured at 37 °C before and after a 1 min DNA denaturation step at 95 °C (in blue shaded region) are depicted within the pink shaded regions.

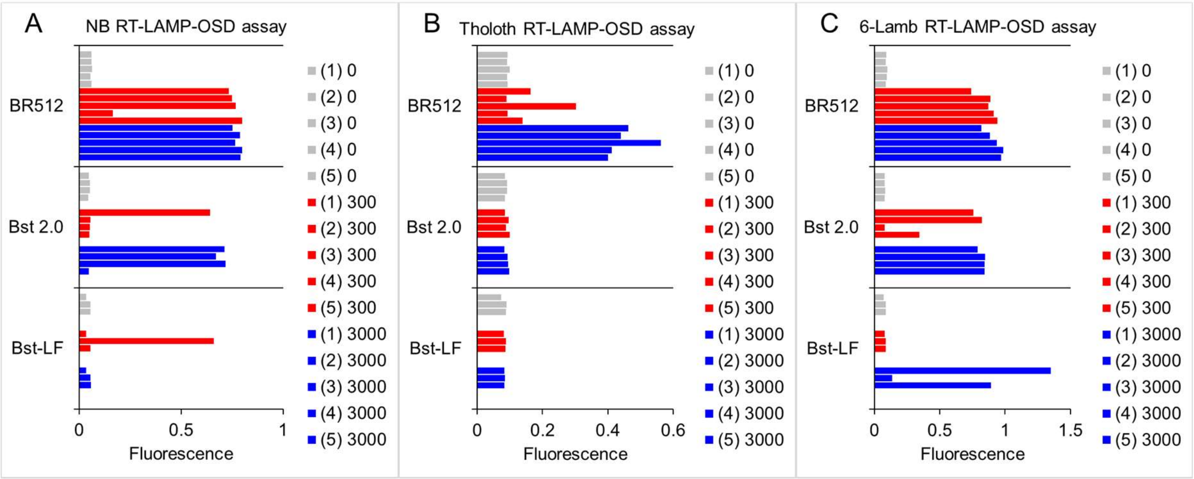


**Supplementary Figure 4. Comparison of Br512, Bst-LF, and Bst 2.0 in RT-LAMP-OSD assays for SARS-CoV-2 genomic RNA.** Three SARS-CoV-2-specific RT-LAMP-OSD assays, NB (panel A), Tholoth (panel B), and 6-Lamb (panel C), were carried out with 20 pm of in-house purified Bst-LF, 16 units of commercially sourced Bst 2.0, or 20 pm of Br512 in 1X G6D, 1X isothermal, and 1X G6D reaction buffers, respectively. OSD fluorescence measured at assay endpoint in reactions seeded with 3,000 (blue bars), 300 (red bars), or 0 (gray bars) copies of SARS-CoV-2 viral genomic RNA templates are depicted. Assay replicates in each panel are numbered 1 through 5. The real-time amplification kinetics of each reaction are detailed in **Supplementary Figure 5**.

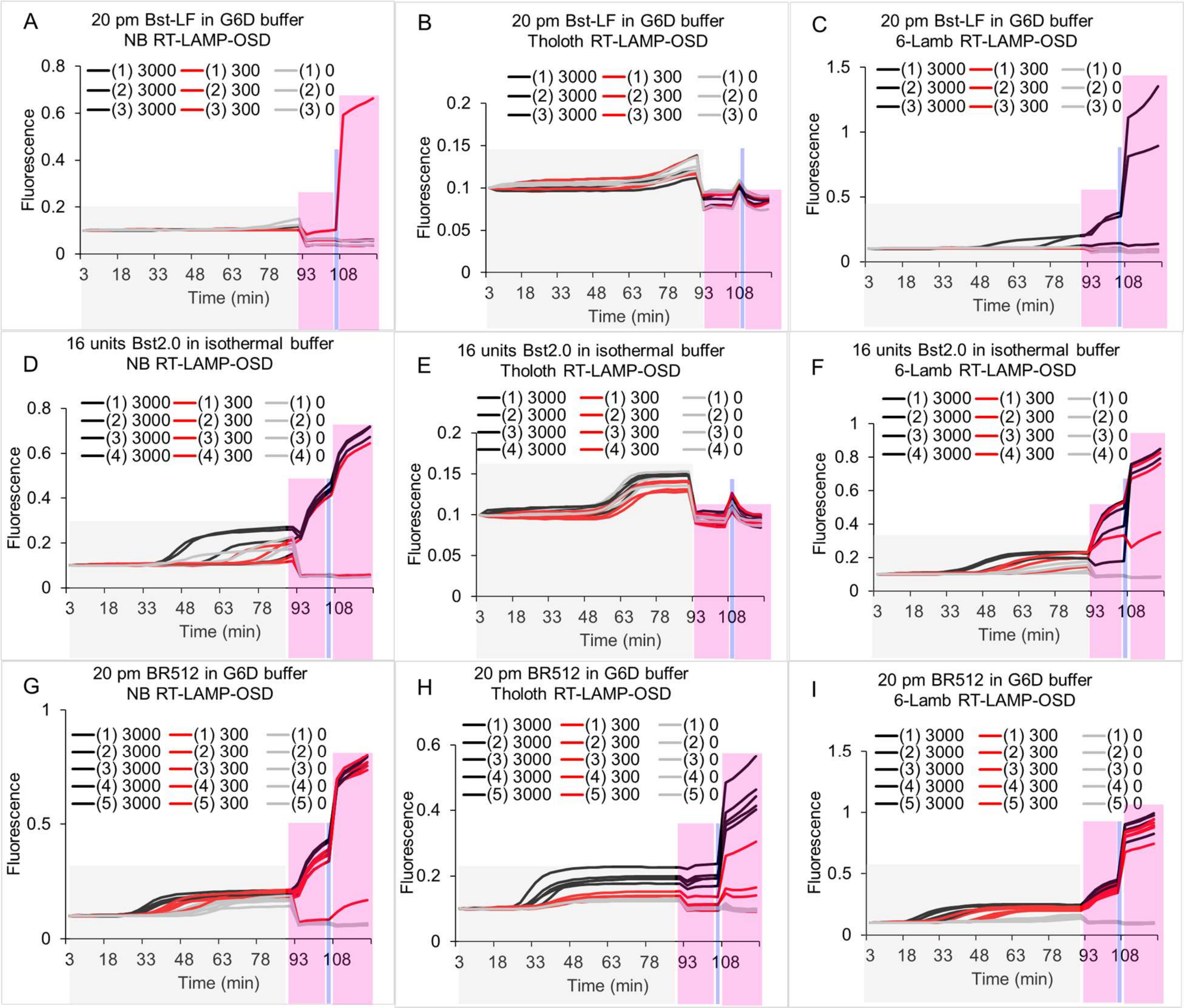


**Supplementary Figure 5. Comparison of Br512, Bst-LF, and Bst 2.0 in RT-LAMP-OSD assays for SARS-CoV-2 genomic RNA.** Three SARS-CoV-2-specific RT-LAMP-OSD assays, NB (panels A, D, and G), Tholoth (panels B, E, and H), and 6-Lamb (panels C, F, and I), were operated using 20 pm of in-house purified Bst-LF (panels A, B, and C), 16 units of commercially sourced Bst 2.0 (panels D, E, and F), or 20 pm of Br512 (panels G, H, and I) in indicated reaction buffers. Amplification kinetics at 65 °C observed in real-time by measuring OSD fluorescence in reactions seeded with 3,000 (black traces), 300 (red traces), or 0 (gray traces) copies of SARS-CoV-2 viral genomic RNA templates are depicted within gray shaded boxes. Post-amplification OSD signal measured at 37 °C before and after a 1 min DNA denaturation step at 95 °C (in blue shaded region) are depicted within the pink shaded regions. Assay replicates in each panel are numbered 1 through 5.

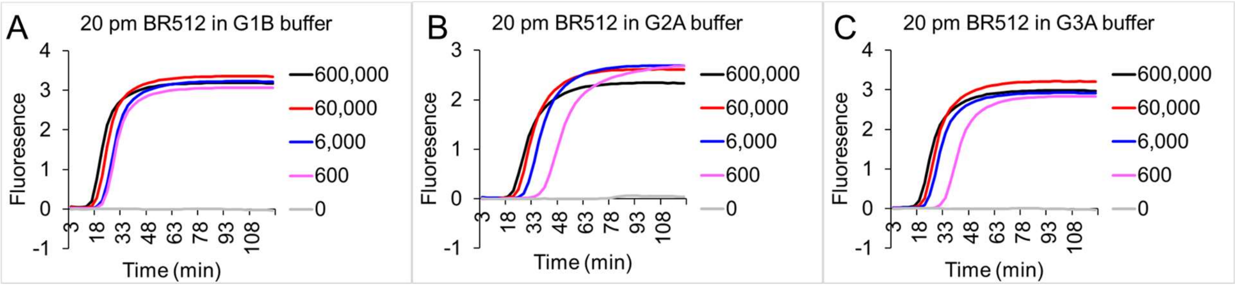


**Supplementary Figure 6. Comparison of Br512 activity in different LAMP-OSD assay buffers.** LAMP-OSD assays with the human *GAPDH* gene were carried out with Br512 in the indicated reaction buffers. Amplification curves were observed in real-time by measuring OSD fluorescence at 65 °C in reactions seeded with 600,000 (black traces), 60,000 (red traces), 6,000 (blue traces), 600 (pink traces), and 0 (gray traces) copies of *GAPDH* plasmid templates.

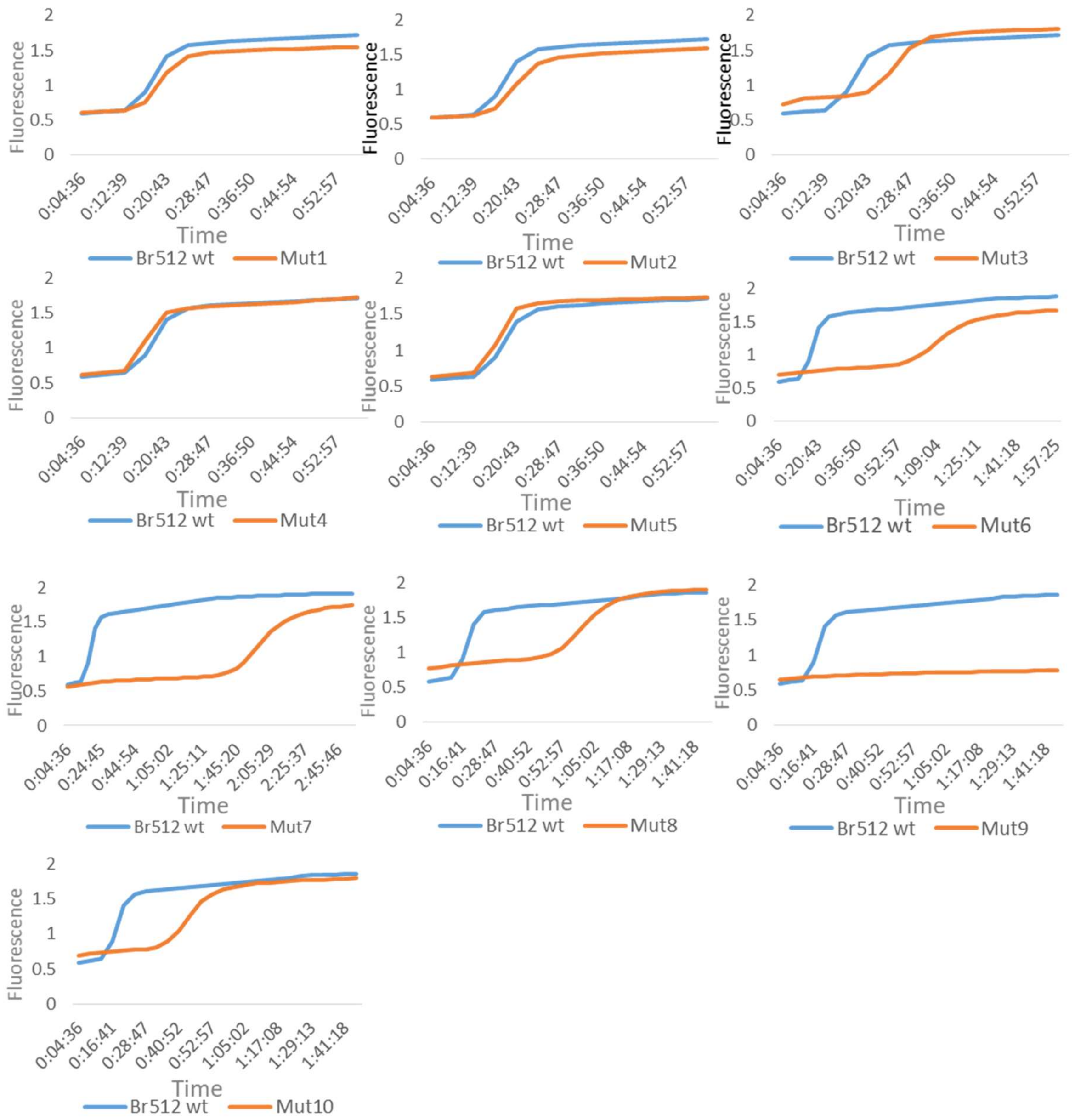


**Supplementary Figure 7. Initial evaluation of computationally predicted substitutions on Br512 (Bst-LF) activity.**

LAMP assays were carried out with a 20 pg ($6x10^7$ copies) of *GAPDH* DNA template to assess the effect of the individual mutations suggested by Mutcompute on Br512 activity. Amplification was observed by EvaGreen dye fluorescence change (Y-axis) over time of incubation (X-axis) at 65°C. Blue traces indicates Br512 wild type and burnt orange traces are individual mutations (Mut1-Mut10).

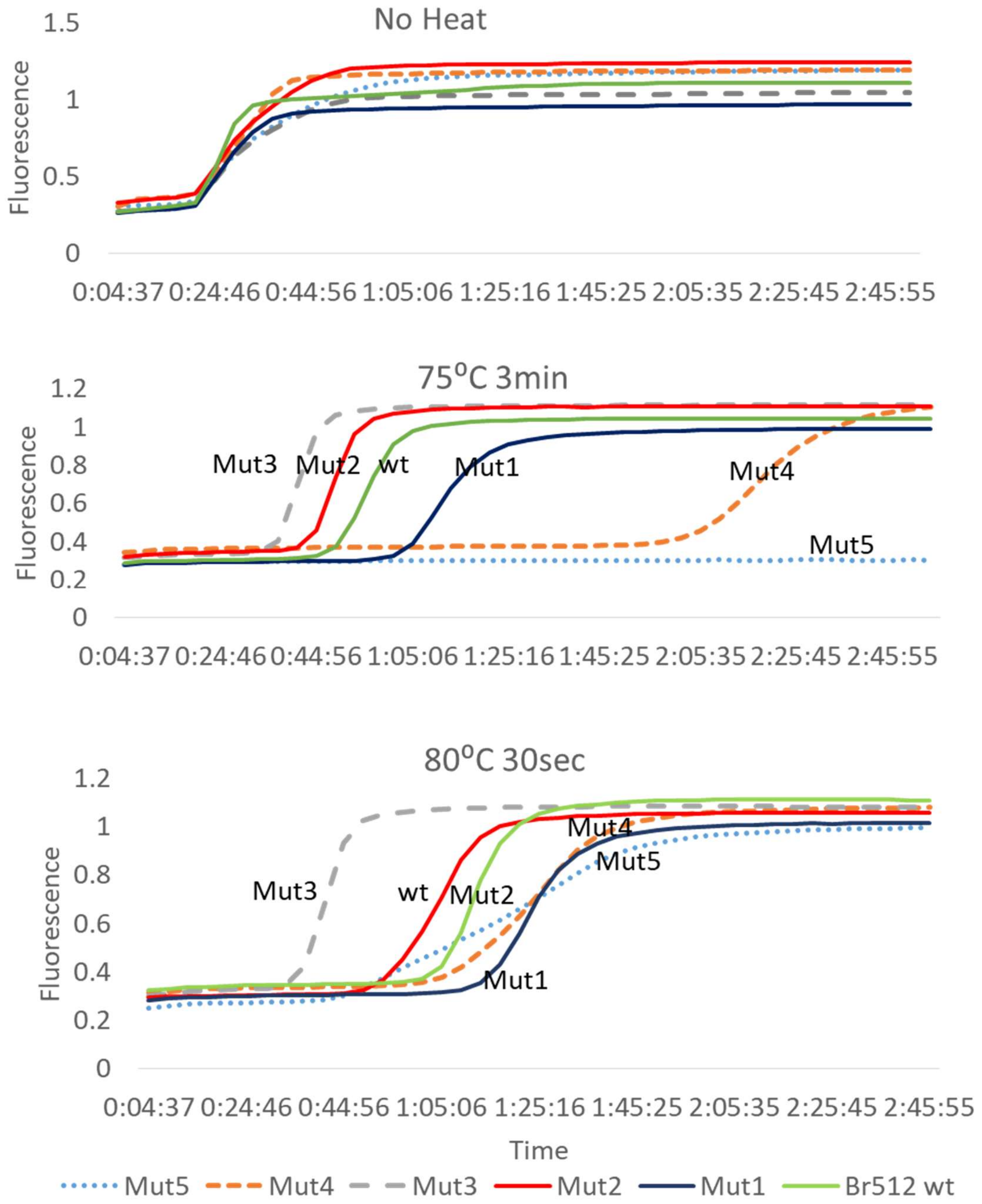


**Supplementary Figure 8. Heat challenge LAMP assay with computationally predicted single amino acid substitutions.** LAMP assays assembled with wildtype (wt) or Mutcompute calculated Br512 variants (Mut1 to 5) were subjected to indicated thermal challenges (top panel: no thermal challenge; middle panel: 3 min at 75°C; lower panel: 30 sec at 80°C) prior to real time measurement of DNA amplification during continuous incubation at 65°C. Amplification kinetics was determined by measuring EvaGreen fluorescence (Y-axis) over incubation time (X-axis; hh:mm:ss). Green: Br512 wt (wild type), Dark blue: Mut1, Red: Mut2, Dotted gray: Mut3, Dotted orange: Mut4, Dotted blue: Mut5

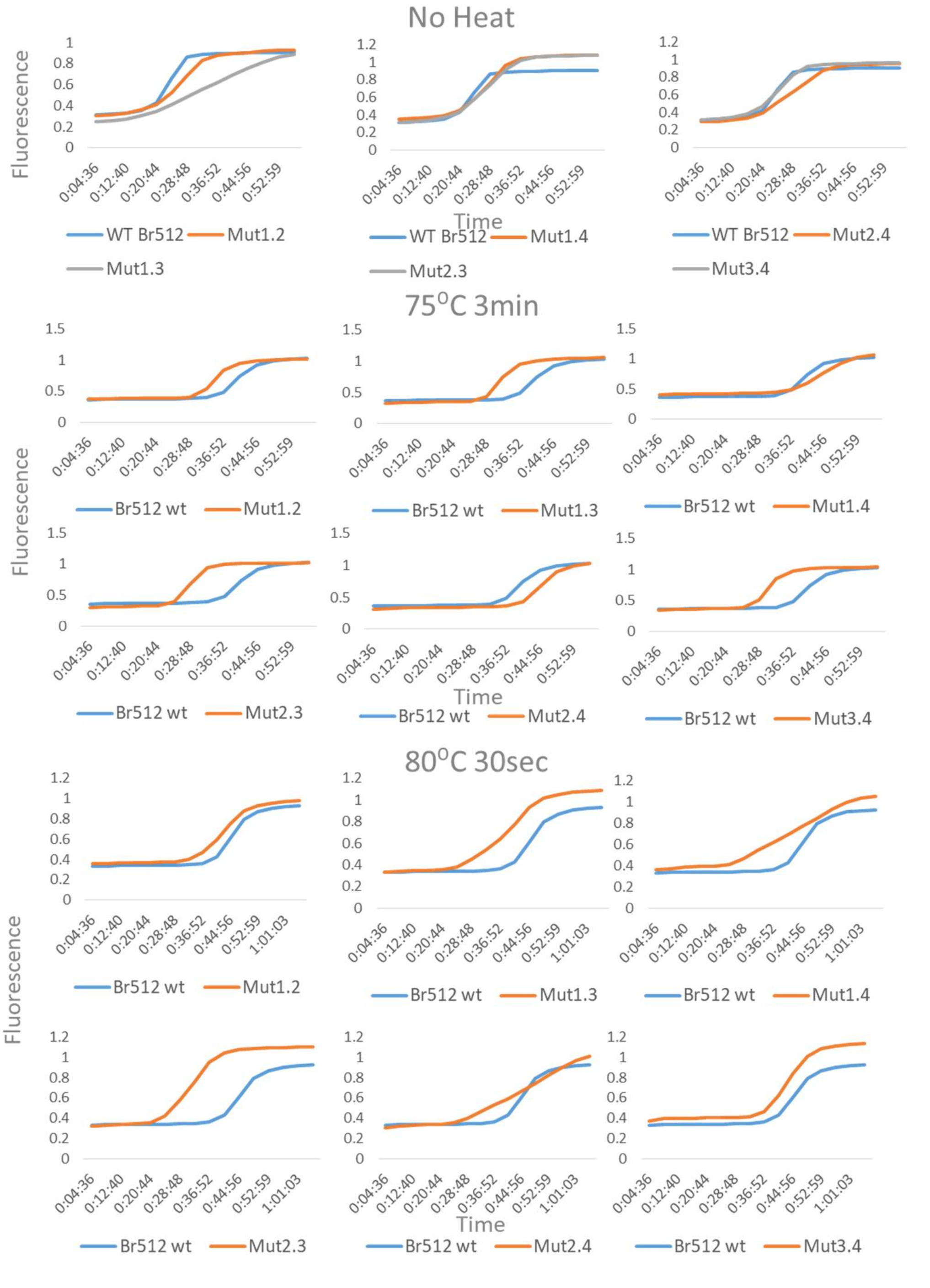


**Supplementary Figure 9. Heat challenge LAMP assay with double mutation Br512 variants.** Activities of wild type (blue traces) and the various double mutant Mutcompute Br512 variants (orange traces) were compared in identical LAMP assays containing 20 pg ($6x10^7$ copies) of *GAPDH* DNA templates that were subjected to indicated thermal challenges (top panel: no thermal challenge; middle panel: 3 min at 75°C; lower panel: 30 sec at 80°C) prior to real time measurement of DNA amplification at 65°C. Representative amplification curves determined by measuring EvaGreen fluorescence (Y-axis) over incubation time (X-axis; hh:mm:ss) are depicted.

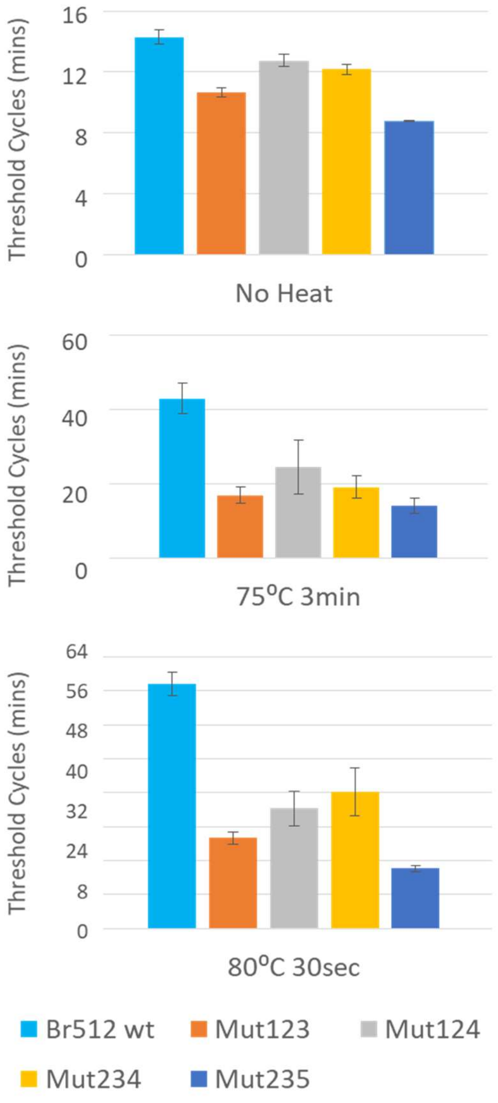


**Supplementary Figure 10. Threshold cycle (Ct) analysis of triple Mutcompute variants.** *GAPDH* LAMP assay results shown in Figure 3 b-d were further quantified with Ct values in minutes. Threshold cycles for amplification of 20 pg ($6x10^7$ copies) *GAPDH* DNA templates were calculated using the Lightcycler96 software (Roche). Lower Ct indicates faster amplification. Upper panel: Ct values for No Heat LAMP, Middle panel: Ct values for 75ºC 3min heat challenge LAMP, Lower panel: Ct values for 80ºC 30sec heat challenge LAMP. , Error bar=S.D., n=2 for No Heat, n=3 for 75ºC and 80ºC heat challenge LAMP (Y-axis: Ct in minutes).

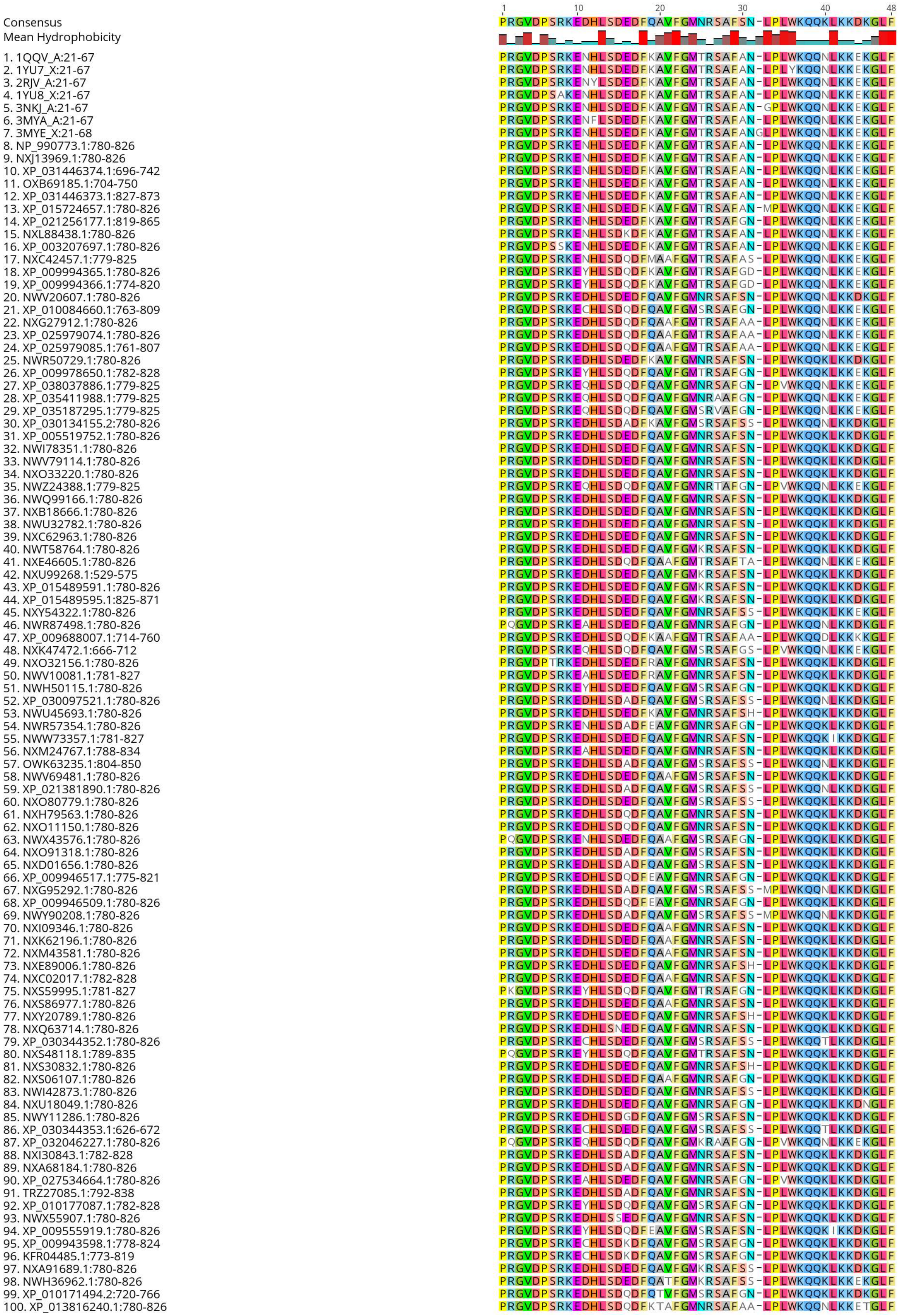
1
10
20
30
40
48
Consensus
PRGVDPSRKEDHLSDEDFQAVFGMNRSAFSN-LPLWKQQKLKKDKGLF
Mean Hydrophobicity
1. 1QQV_A:21-67
2. 1YU7_X:21-67
3. 2RJV_A:21-67
4. 1YU8_X:21-67
5. 3NKJ_A:21-67
6. 3MYA_A:21-67
7. 3MYE_X:21-68
8. NP_990773.1:780-826
9. NXJ13969.1:780-826
10. XP_031446374.1:696-742
11. OXB69185.1:704-750
12. XP_031446373.1:827-873
13. XP_015724657.1:780-826
14. XP_021256177.1:819-865
15. NXL88438.1:780-826
16. XP_003207697.1:780-826
17. NXC42457.1:779-825
18. XP_009994365.1:780-826
19. XP_009994366.1:774-820
20. NWV20607.1:780-826
21. XP_010084660.1:763-809
22. NXG27912.1:780-826
23. XP_025979074.1:780-826
24. XP_025979085.1:761-807
25. NWR50729.1:780-826
26. XP_009978650.1:782-828
27. XP_038037886.1:779-825
28. XP_035411988.1:779-825
29. XP_035187295.1:779-825
30. XP_030134155.2:780-826
31. XP_005519752.1:780-826
32. NWI78351.1:780-826
33. NWV79114.1:780-826
34. NXO33220.1:780-826
35. NWZ24388.1:779-825
36. NWQ99166.1:780-826
37. NXB18666.1:780-826
38. NWU32782.1:780-826
39. NXC62963.1:780-826
40. NWT58764.1:780-826
41. NXE46605.1:780-826
42. NXU99268.1:529-575
43. XP_015489591.1:780-826
44. XP_015489595.1:825-871
45. NXY54322.1:780-826
46. NWR87498.1:780-826
47. XP_009688007.1:714-760
48. NXK47472.1:666-712
49. NXO32156.1:780-826
50. NWV10081.1:781-827
51. NWH50115.1:780-826
52. XP_030097521.1:780-826
53. NWU45693.1:780-826
54. NWR57354.1:780-826
55. NWW73357.1:781-827
56. NXM24767.1:788-834
57. OWK63235.1:804-850
58. NWV69481.1:780-826
59. XP_021381890.1:780-826
60. NXO80779.1:780-826
61. NXH79563.1:780-826
62. NXO11150.1:780-826
63. NWX43576.1:780-826
64. NXO91318.1:780-826
65. NXD01656.1:780-826
66. XP_009946517.1:775-821
67. NXG95292.1:780-826
68. XP_009946509.1:780-826
69. NWY90208.1:780-826
70. NXI09346.1:780-826
71. NXK62196.1:780-826
72. NXM43581.1:780-826
73. NXE89006.1:780-826
74. NXC02017.1:782-828
75. NXS59995.1:781-827
76. NXS86977.1:780-826
77. NXY20789.1:780-826
78. NXQ63714.1:780-826
79. XP_030344352.1:780-826
80. NXS48118.1:789-835
81. NXS30832.1:780-826
82. NXS06107.1:780-826
83. NWI42873.1:780-826
84. NXU18049.1:780-826
85. NWY11286.1:780-826
86. XP_030344353.1:626-672
87. XP_032046227.1:780-826
88. NXI30843.1:782-828
89. NXA68184.1:780-826
90. XP_027534664.1:780-826
91. TRZ27085.1:792-838
92. XP_010177087.1:782-828
93. NWX55907.1:780-826
94. XP_009555919.1:780-826
95. XP_009943598.1:778-824
96. KFR04485.1:773-819
97. NXA91689.1:780-826
98. NWH36962.1:780-826
99. XP_010171494.2:720-766
100. XP_013816240.1:780-826

**Supplementary Figure 11. Alignment of vHP47 sequence and its conserved amino acids among its orthologues.**

Amino acid sequence of villin headpiece vHP47 was blasted at https://blast.ncbi.nlm.nih.gov/Blast.cgi with blastp (protein-protein BLAST) algorithm. Top 100 hit sequences were compared using NCBI Multiple Sequences Alignment Viewer. Amino acids that are identical to consensus sequence were highlighted. Mean hydrophobicity is shown as a bar graph. Top row: consensus amino acid sequence.

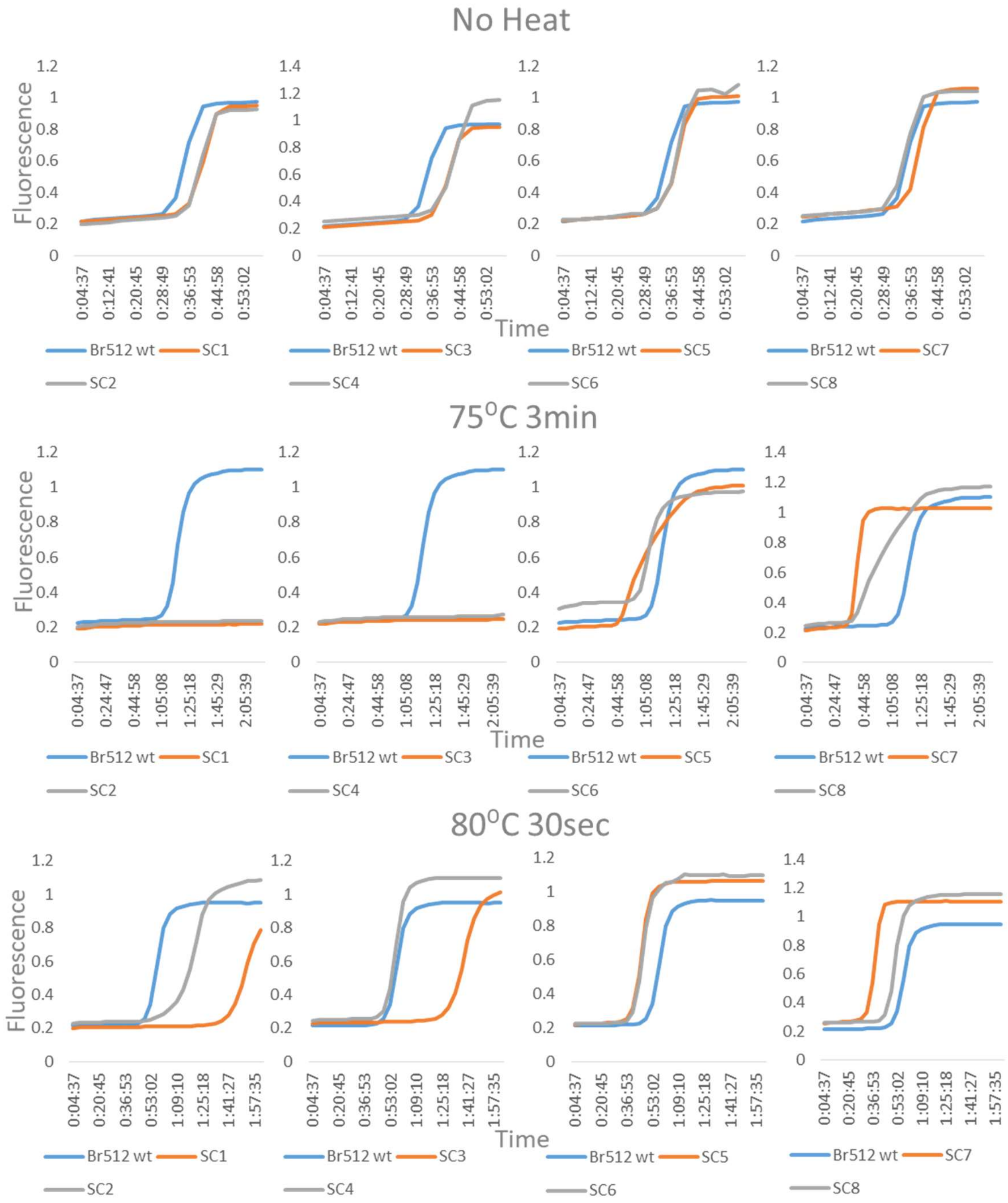


**Supplementary Figure 12. Heat challenge of single mutation supercharged Br512 variants.** Identical LAMP assays assembled using wild type (blue) or mutant (orange and gray) Br512 were heat challenged at either 75°C for 3min or 80°C for 30sec prior to determining amplification kinetics at 65 °C Representative amplification curves determined by measuring EvaGreen fluorescence (Y-axis) over incubation time (X-axis; hh:mm:ss) are depicted.

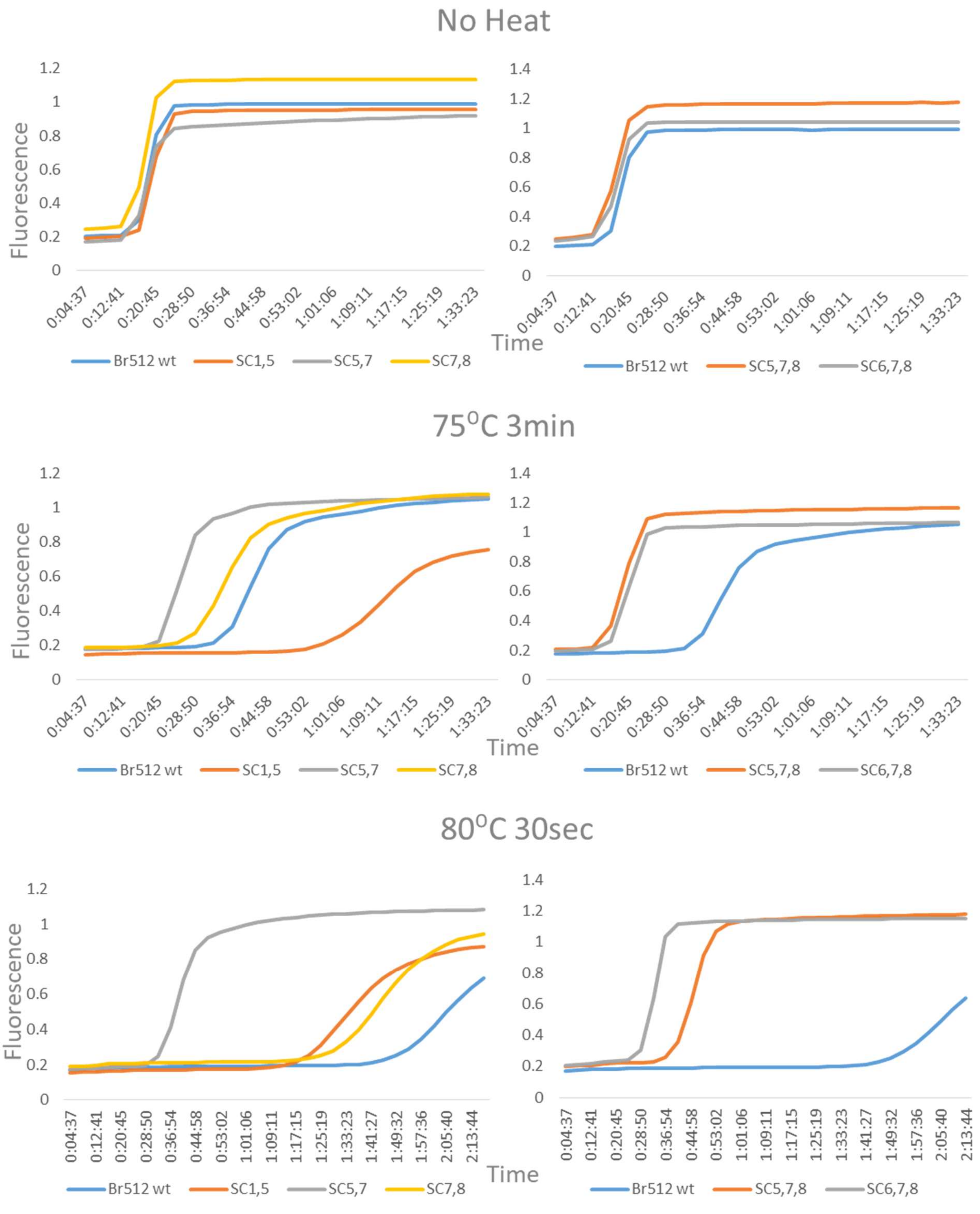


**Supplementary Figure 13. Heat challenge of supercharged double and triple mutation Br512 variants.**

Identical *GAPDH* LAMP assays assembled using either double mutation variants (Left panels) or triple mutation variants (right panels) were subjected to the indicated heat challenges prior to measuring amplification kinetics at 65 °C. Representative amplification curves determined by measuring EvaGreen fluorescence (Y-axis) over incubation time (X-axis; hh:mm:ss) are depicted as blue (wild type Br512) or burnt orange, yellow and gray (indicated double and triple mutants) traces.

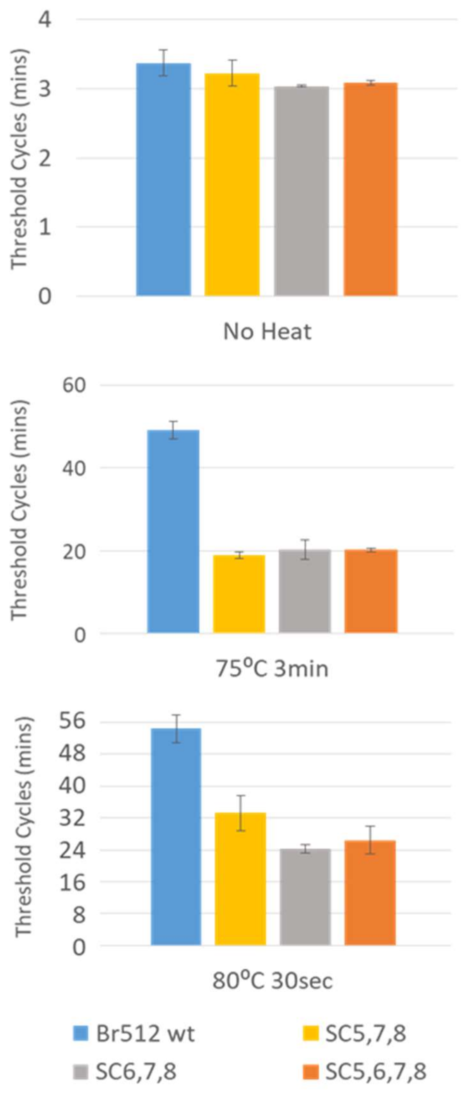


**Supplementary Figure 14. Threshold cycle ($C_t$) analysis of triple and quadruple supercharged vHP47 variants.** LAMP assay results shown in Figure 4c were further quantified by $C_t$ values in minutes. Threshold cycles for amplification of 20 pg ($6x10^7$ copies) *GAPDH* DNA templates were calculated by Lightcycler96 software. Error bar=S.D., n=2 for No Heat, n=3 for 75ºC and 80ºC heat challenge LAMP. (Y-axis: Ct in minutes).

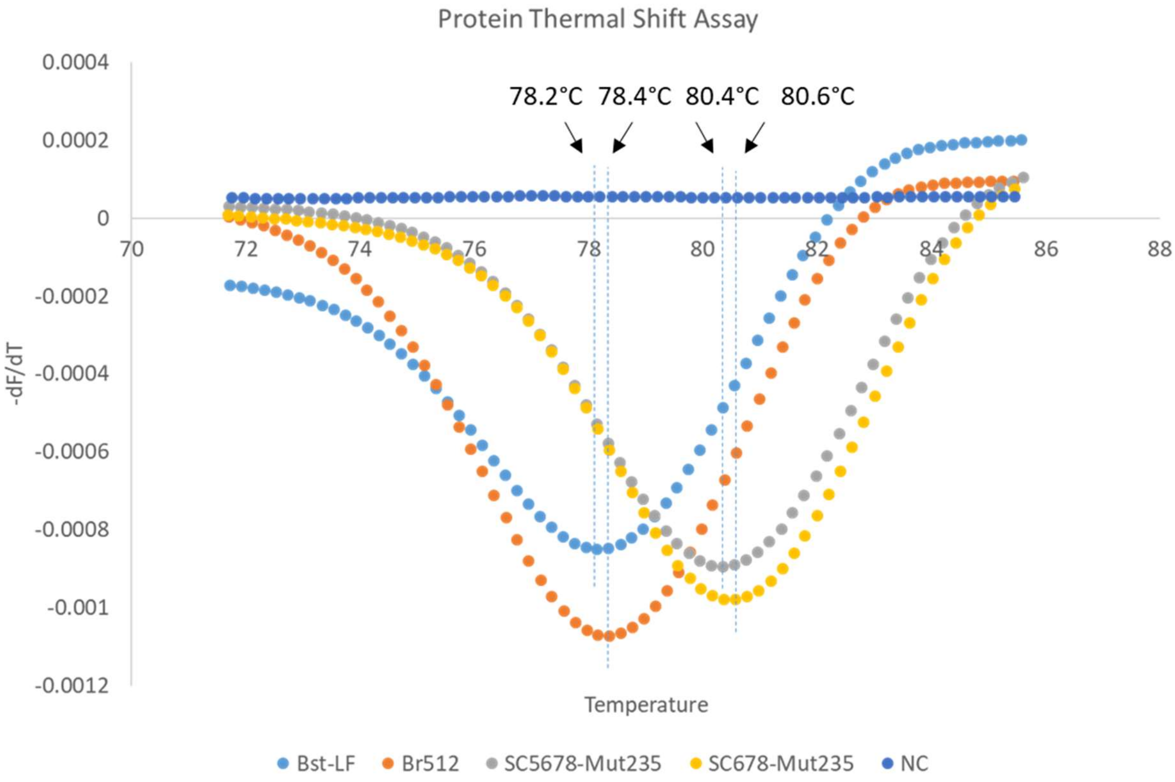


**Supplementary Figure 15. Protein Thermal Shift Assay for wildtype and mutant Br512 variants.**

Same amount (40µg) of parental (Bst-LF) and engineered enzyme variants were analyzed using Protein Thermal Shift$^{TM}$ (Thermo Fisher; Catalog Number: 4461146), a dye-based protein thermal shift assay, according to the manufacturer's instructions. The enzymes were incubated in a Lightcyler 96 (Roche) real-time PCR machine programmed to ramp temperature from 37 °C to 95 °C at the rate of 0.1 °C/sec while continuously measuring changes in red fluorescence. Melt curves generated by plotting change in fluorescence (dF) as a function of changing temperature (dT) are depicted.  NC: No-protein Control.

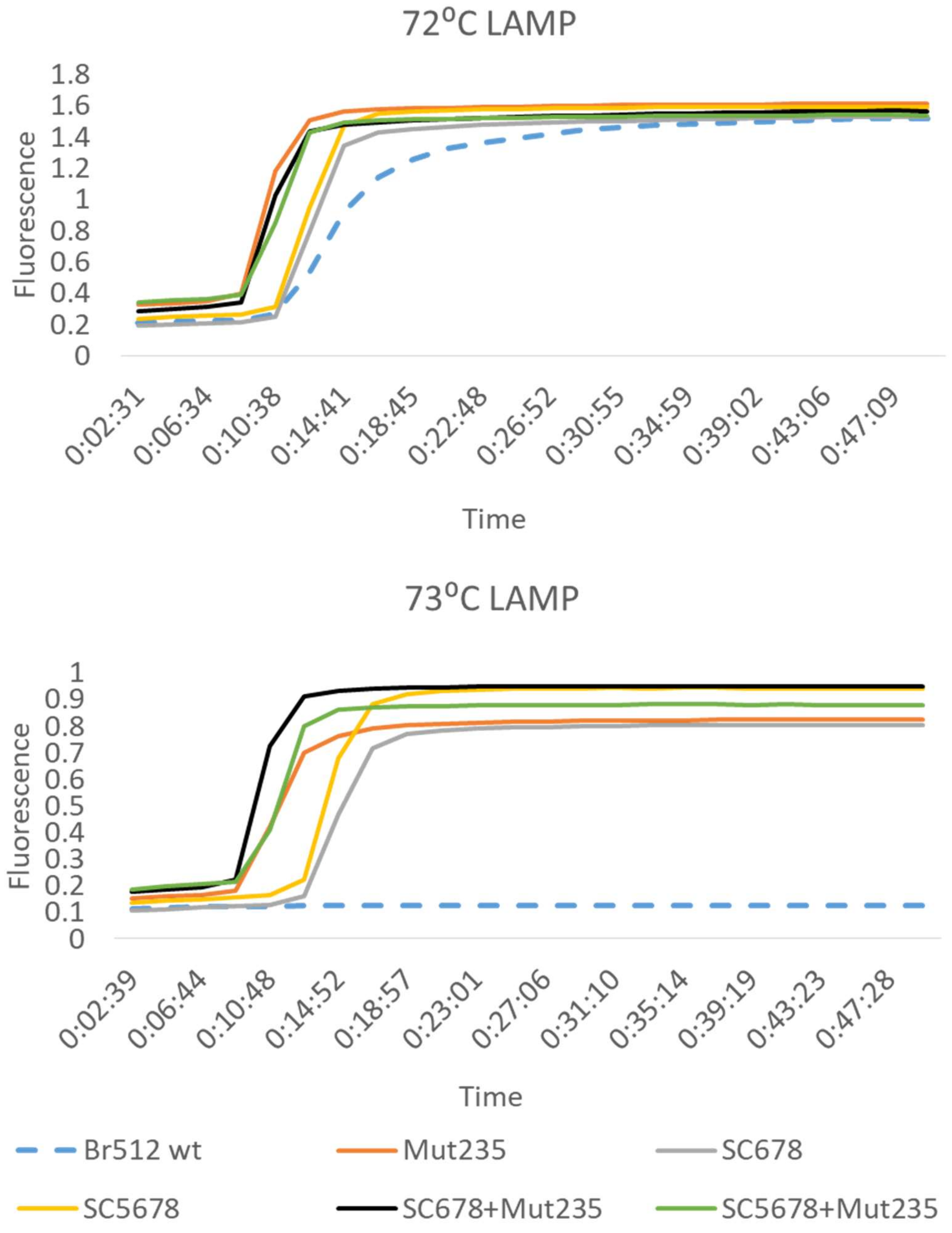


**Supplementary Figure 16. High temperature *GAPDH* LAMP at 72ºC and 73ºC.**
Identical *GAPDH* LAMP assays were assembled using equal amounts of either wildtype or indicated mutant Br512 variants and DNA amplification kinetics at 72 ºC or 73 ºC was determined by measuring changes in EvaGreen fluorescence. Representative amplification curves showing change in fluorescence (Y-axis) over time (X- axis; hh:mm:ss) are depicted as dotted blue dotted (Br512 wild type), burnt orange (Mut235), gray (SC678), black (SC678+Mut235), and green (SC5678+Mut235) traces.

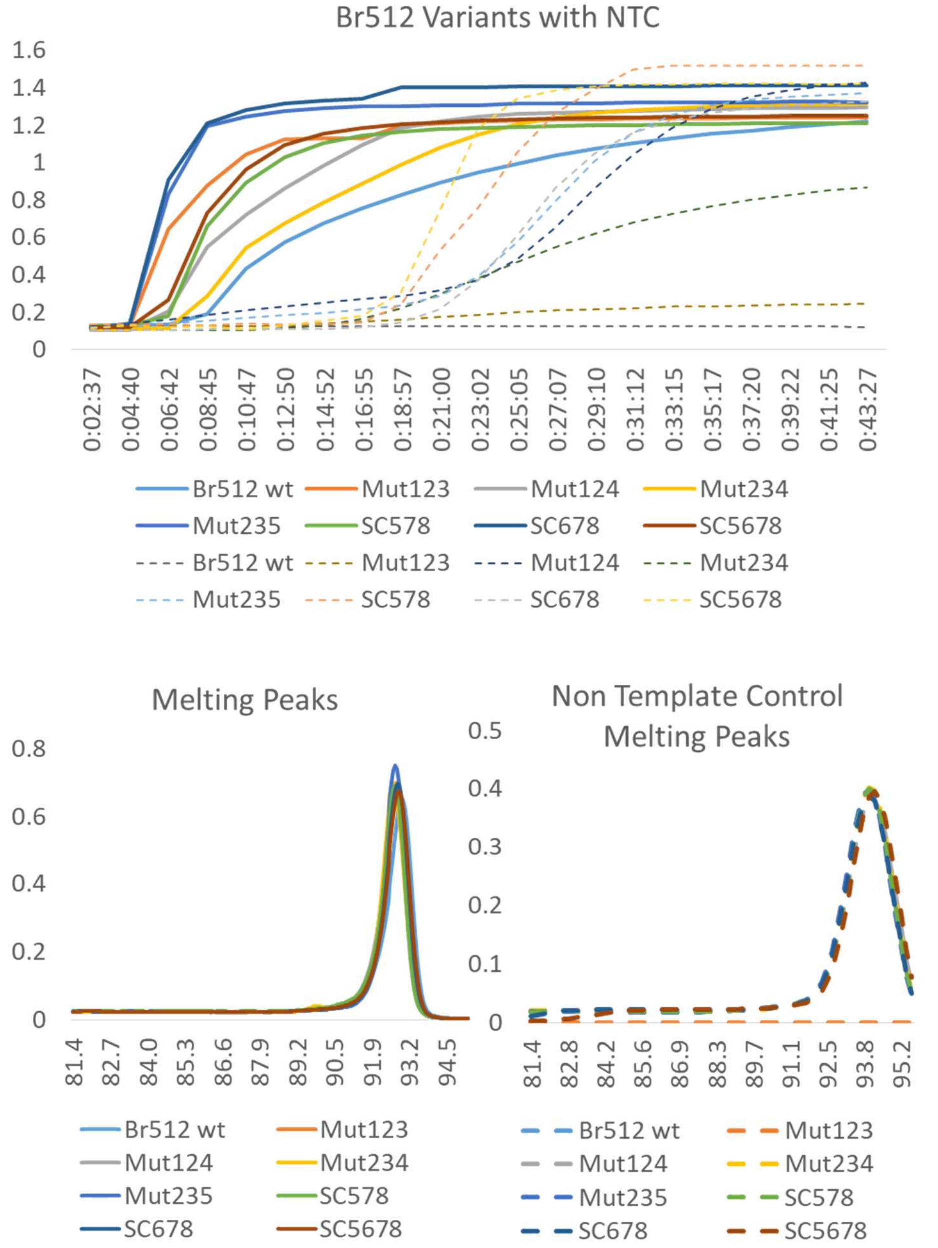


**Supplementary Figure 17. Non-template controls in the thermal challenge LAMP assays. (Upper panel)** Amplification curves observed in *GAPDH* LAMP assays containing indicated enzymes and 20 pg (6x$10^7$ copies) *GAPDH* DNA templates are depicted as solid traces while corresponding NTC assays without specific templates are plotted as dotted traces. NTC either present delayed amplification or no amplification. (Y-axis: Fluorescence intensity X-axis: time of incubation) **(Lower panels)** Melting curve analysis of amplicons generated in LAMP assays with 20 pg (6x$10^7$ copies) *GAPDH* DNA templates (left panel) or NTC LAMP assays without templates (right panel). (Y-axis: -dF/dT: delta Fluorescence/delta temperature X-axis: temperature in °C)

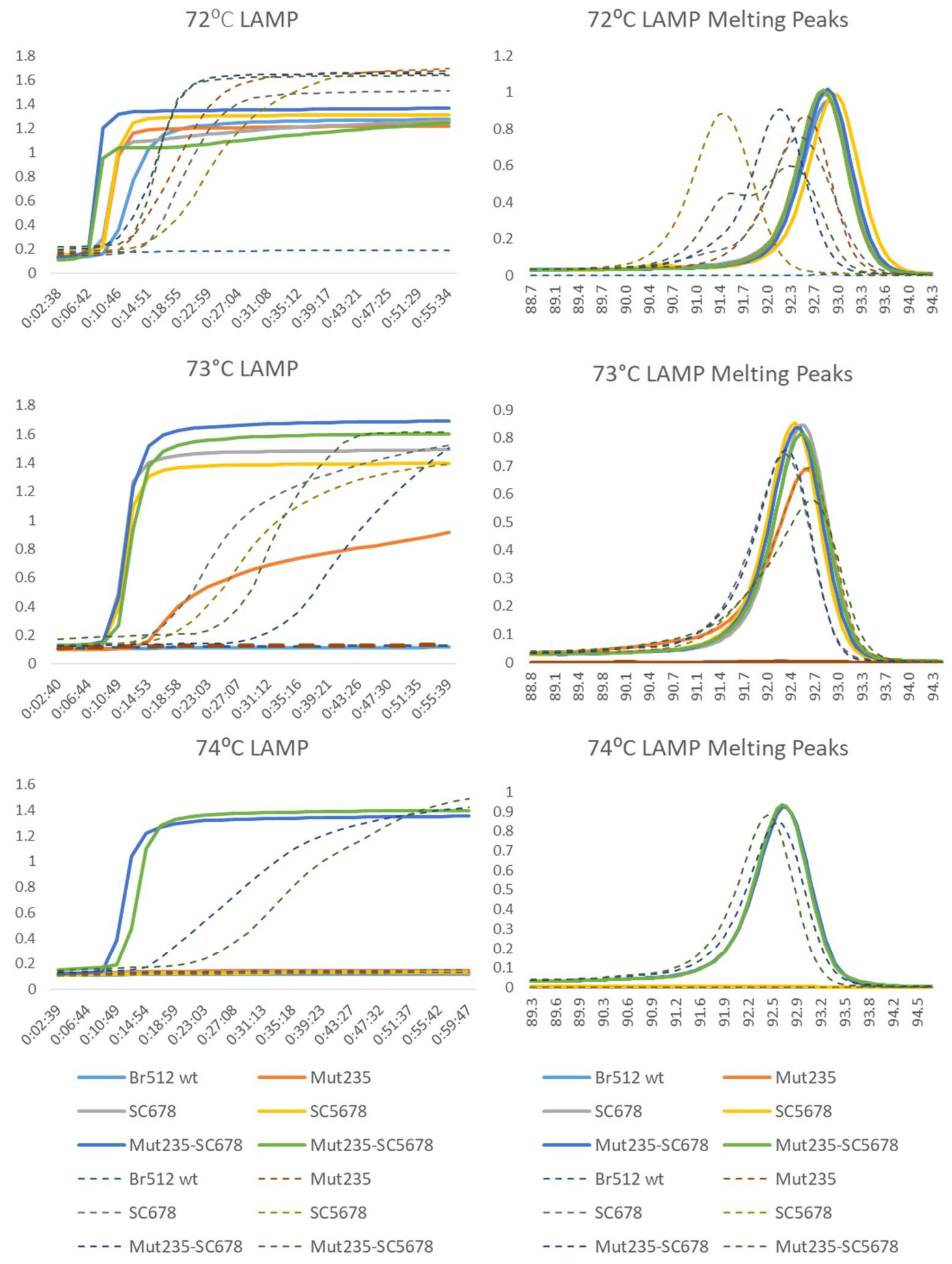


**Supplemental Figure 18. Non-template controls in the high temperature LAMP assays.** Representative amplification results from various high temperature *GAPDH* LAMP assays were plotted. Solid lines indicate amplification curves from reactions seeded with 20 pg ($6x10^7$ copies) of *GAPDH* DNA templates and dotted lines indicate corresponding non-template controls (NTC). Left three panels show amplifications curve (Y-axis: Fluorescence intensity X-axis: time of incubation). Right three panels show melting peaks (Y-axis: -dF/dT: delta Fluorescence/delta temperature X-axis: temperature in °C)

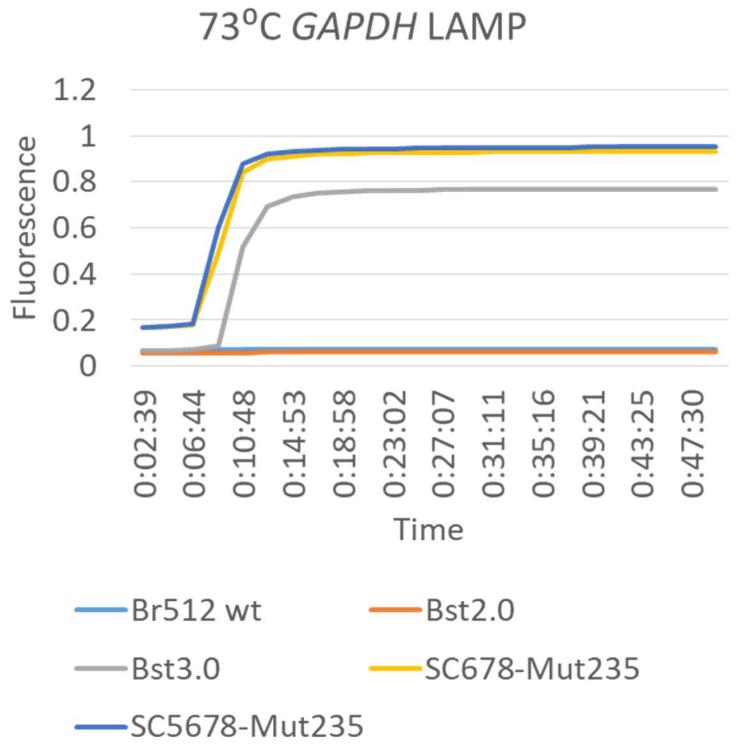


**Supplementary Figure 19. Comparison of supercharged vHP47-Mut Br512 variants, Bst2.0, and Bst3.0 enzymes in 73°C *GAPDH* LAMP.**

*GAPDH* DNA LAMP assays containing either 16 units of Bst3.0 (NEB) or 16 units of Bst2.0 (NEB) in 1X Isothermal II buffer (NEB) with 8mM $MgSO_4$ or containing 20 pmol of wildtype or mutant Br512 variants in the same reaction buffer were incubated at 73 °C and DNA amplification was evaluated by measuring EvaGreen fluorescence. Representative amplification curves showing change in fluorescence (Y-axis) over time (X-axis) are depicted.

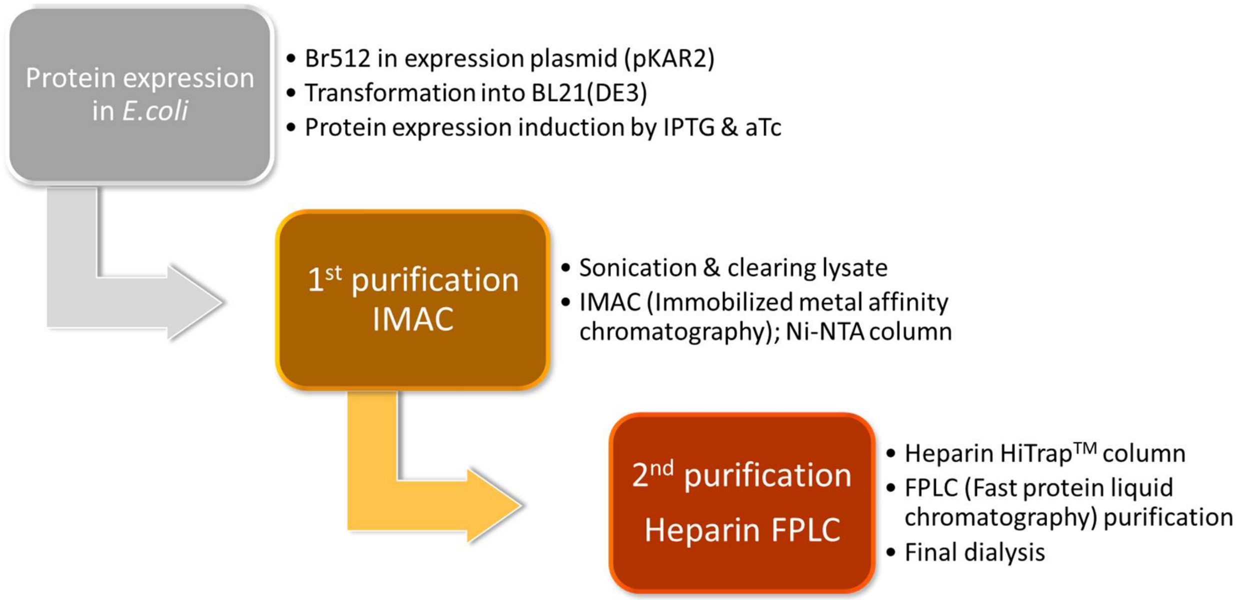


**Supplementary Figure 20. A flowchart of simple two-step Br512 purification.** Simple two-step purification procedures are shown in the flowchart. *E. coli* BL21(DE3) cell expressed Br512 was initially purified with Ni-NTA based immobilized metal affinity chromatography (IMAC), and further purified with heparin column based FPLC. A detailed purification protocol is described in the materials and methods section.

| Supplementary Table 1. Oligonucleotide and template sequences used in the study | | |
|---|---|---|
| **Name** | **Sequence** | **Use** |
| gapdLAMP.F3 | GCCACCCAGAAGACTGTG | gapd LAMP-OSD |
| gapdLAMP.B3 | TGGCAGGTTTTTCTAGACGG | |
| gapdLAMP.FIP | CGCCAGTAGAGGCAGGGATGAGGGAAACTGTGGCGTGAT | |
| gapdLAMP.BIP | GGTCATCCCTGAGCTGAACGGTCAGGTCCACCACTGACAC | |
| gapdLAMP.LR | TGTTCTGGAGAGCCCCGCGGCC | |
| gapdOSD.F | /56-FAM/CTCACTGGCATGGCCTTCCGTGTCCCCACTGCCAAC/3InvdT/ | |
| gapdOSD.Q | GGACACGGAAGGCCATGCCAGTGAG/3IABkFQ/ | |
| gapd template | CTAGTAACGGCCGCCAGTGTGCTGGAATTCCCACAGTCCATGCCATCACTGCCACCCAGAAGACTGTGGATGGCCCCTCCGGGAAACTGTGGCGTGATGGCCGCGGGGCTCTCCAGAACATCATCCCTGCCTCTACTGGCGCTGCCAAGGCTGTGGGCAAGGTCATCCCTGAGCTGAACGGGAAGCTCACTGGCATGGCCTTCCGTGTCCCCACTGCCAACGTGTCAGTGGTGGACCTGACCTGCCGTCTAGAAAAACCTGCCAAATATGATGACATCAAGAAGGTGGTGAAGCAGGCGTCGGAGGGCCCCCTCAAGGGCATCCTGGGCTACACTGAGCACCAGGTGGTCTCCTCTGACTTCAACAGCGACACCCACTCCTCCACCTTTGACGCTGGGGCTGGCATTGCCCTCAACGACCACTTTGTCAAGCTCATTTCCTGGAATTCTGCAGATATCCATCACACTGGCGGCCGCTCGAGC | |
| NB-F3 | ACCGAAGAGCTACCAGACG | NB RT-LAMP-OSD |
| NB-B3 | TGCAGCATTGTTAGCAGGAT | |
| NB-FIP | TCTGGCCCAGTTCCTAGGTAGTTCGTGGTGGTGACGGTAA | |
| NB-BIP | AGACGGCATCATATGGGTTGCACGGGTGCCAATGTGATCT | |
| NB-LB | ACTGAGGGAGCCTTGAATACA | |
| NB-OSD-FAM | /56-FAM/CCGAATGAAAGATCTCAGTCCAAGATGGTATTTCT/3InvdT/ | |
| NB-OSD-Q | TCTTGGACTGAGATCTTTCATTCGG/3IABkFQ/ | |
| Lamb-F3 | TCCAGATGAGGATGAAGAAGA | 6-Lamb RT-LAMP-OSD |
| Lamb-B3 | AGTCTGAACAACTGGTGTAAG | |
| Lamb-FIP | AGAGCAGCAGAAGTGGCACAGGTGATTGTGAAGAAGAAGAG | |
| Lamb-BIP | TCAACCTGAAGAAGAGCAAGAACTGATTGTCCTCACTGCC | |
| Lamb-LF | CTCATATTGAGTTGATGGCTCA | |
| Lamb-LB | ACAAACTGTTGGTCAACAAGAC | |
| Lamb-OSD-FAM | GTATGGTACTGAAGATGATTACCAAGGTAAACCTTTGGAATTTGGAC/36-FAM/ | |
| Lamb-OSD-Q | /5IABkFQ/GTCCAAATTCCAAAGGTTTACCTTGGTAATCATCTC/3InvdT/ | |
| Tholoth-F3 | TGCTTCAGTCAGCTGATG | Tholoth RT-LAMP-OSD |
| Tholoth-B3 | TTAAATTGTCATCTTCGTCCTT | |
| Tholoth-FIP | TCAGTACTAGTGCCTGTGCCCACAATCGTTTTAAACGGGT | |
| Tholoth-BIP | TCGTATACAGGGCTTTTGACATCTATCTTGGAAGCGACAACAA | |
| Tholoth-LB | GTAGCTGGTTTTGCTAAATTCC | |
| Tholoth-OSD-FAM | /56-FAM/ACAGGTGTAAGTGCAGCCCGTCTTACACCGTGC/3InvdT/ | |
| Tholoth-OSD-Q | GACGGGCTGCACTTACACCTGT/3IABkFQ/ | |
| HP47 R31E F | CATCCGGAAGGTGTTGACCCGAGC | SC1 |
| HP47 R31E R | GTGGTGATGACCGTGATGATGGTGCATATG | |
| HP47 G32D F | CCGCGTGATGTTGACCCGAGCCGTAAGGAGAAC | SC2 |
| HP47 G32D R | ATGGTGGTGATGACCGTGATGATGGTGCATATG | |
| HP47 R37E F | CCGAGCGAAAAGGAGAACCACCTGTCTGAC | SC3 |
| HP47 R37E R | GTCAACACCACGCGGATGGTGGTGATGACC | |
| HP47 K38D F | AGCCGTGATGAGAACCACCTGTCTGACGAAGAC | SC4 |
| HP47 K38D R | CGGGTCAACACCACGCGGATGGTGGTGATG | |
| HP47 A49K F | TTCAAGAAAGTGTTCGGTATGACCCGTTCTGCG | SC5 |
| HP47 A49K R | GTCTTCGTCAGACAGGTGGTTCTCCTTAC | |
| HP47 N60R F | TTCGCGCGTCTGCCGCTGTGGAAACAACAGAAC | SC6 |
| HP47 N60R R | CGCAGAACGGGTCATACCGAACACCGCCTTG | |
| HP47 E72K F | AAGAAGAAGAAAGGTCTGTTCGGTTCTGGAAGC | SC7 |
| HP47 E72K R | CAGGTTCTGTTGTTTCCACAGCGGCAGGTTC | |
| HP47 N68K F | CAGAAGCTGAAGAAGGAGAAAGGTCTGTTCGGTTCTGGAAGCGCAGCAG | SC8 |

| | | |
|---|---|---|
| HP47 N68K R | TTGTTTCCACAGCGGCAGGTTCGCGAACGC | |
| V191L F | CTGCTTTTAGAGTTAGAACAGCCTC | Mut1 |
| V191L R | GCGGTCCTGCTCGTTGCGAC | |
| T493N F | GATATCAACAGCCGTAACTTCAATGTAC | Mut2 |
| T493N R | AGGAAGGTAGCGACGACGGTG | |
| A552G F | CTTGAGGGCCCCAAGGAAGAGATG | Mut3 |
| A552G R | GATCAATTCGTCATGGACTTGCAGT | |
| R562V F | CTTTGCGTTCTGGTGCCGGAAGTA | Mut4 |
| R562V R | ACGCTCCATCTCTTCCTTGGG | |
| S371D F | GATTACGATCAGATCGAGCTTCGCG | Mut5 |
| S371D R | TGCTGCGAAGATCAGCCAGTCC | |
| N528E F | GACTTACAGGCTCGTCTGAAGGAAG | Mut6 |
| N528E R | GATCATAGCTTTCTTGATGATATCTG | |
| T510F F | ATGAATTTCCCCATCCAGGGGTCAG | Mut7 |
| T510F R | CGCCATGCGTTCGGCGAAG | |
| I304V F | TCTACCTACGTTGAGGGTCTGTTAAAGGTC | Mut8 |
| I304V R | CTGCAGCTTGCCCAGCTGGC | |
| Y303H F | TCTACCCACATTGAGGGTCTGTTAAAGGTC | Mut9 |
| Y303H R | CTGCAGCTTGCCCAGCTGGC | |
| V572A F | GGCCGCGACGCTGCGCGTAC | Mut10 |
| V572A R | TGTTCCATTACTTCCGGCACCAG | |

**Supplementary Table 2. Full sequence of pKAR2-Br512; 6218 bp.** Br512 is highlighted.

Detailed annotation of Br512
4285-4311 : 8x His
4312-4452 : HP47
4453-4476 : 2(GS)3(A)P
4476-6216 : Bst-LF

ATCCTAAGCTTAATTTAGCATAACCCCTTGGGGCCTCTAAACGGGTCTTGAGGGGT
TTTTTGAGCTGAGCTTGGACTCCTGTTGATAGATCCGGCCGGTAATGACCTCAGAA
CTCCATCTGCCTAATGGAGTGATTCTAGCCGGTCGTTACACGTGGAACGGGAACT
GCCAGACATCAAATAAAACAAAAGGCTCAGTCGGAAGACTGGGCCTTTTGTTTTAT
CTGTTGTTTGTCGGTGAACACTCTCCCGGAAAACTCACGTTAAGGGATTTTGGTCA
TGAGATTATCAAAAAGGATCTTCACCTAGATCCTTTTAAACTAGTGAAGTTACCATC
ACGGAAAAAGGTTATGCTGCTTTTAAGACCCACTTTCACATTTAAGTTGTTTTTCTAA
TCCGCAAATGATCAATTCAAGGCCGAATAAGAAGGCTGGCTCTGCACCTTGGTGAT
CAAATAATTCGATAGCTTGTCGTAATAATGGCGGCATACTATCAGTAGTAGGTGTTT
CCCTTTCTTCTTTAGCGACTTGATGCTCTTGATCTTCCAATACGCAACCTAAAGTAA
AATGCCCCACTGCACTGAGTGCATATAATGCATTCTCTAGTGAAAAACCTTGTTGG
CATAAAAAGGCTAATTGATTTTCGAGAGTTTCATACTGTTTTTCTGTAGGCCGTGTA
CCTAAATGTACTTTTGCTCCATCGCGATGACTTAGTAAAGCACATCTAAAACTTTTA
GCGTTATTACGTAAAAAATCTTGCCAGCTTTCCCCTTCTAAAGGGCAAAAGTGAGT
ATGGTGCCTATCTAACATCTCAATGGCTAAGGCGTCGAGCAAAGCCCGCTTATTTT
TTACATGCCAATACAATGTAGGCTGCTCTACACCTAGCTTCTGGGCGAGTTTACGG
GTTGTTAAACCTTCGATTCCGACCTCATTAAGCAGCTCTAATGCGCTGTTAATCACT
TTACTTTTATCTAAACGAGACATCATTAACCTCCTCAAGAGGATCGAATAGTTATTA
CCAATGCTTAATCAGTGAGGCACCTATCTCAGCGATCTGTCTATTTCGTTCATCCAT
AGTTGCCTGACTCCCCGTCGTGTAGATAACTACGATACGGGAGGGCTTACCATCT
GGCCCCAGTGCTGCAATGATACCGCGAGAGCCACGCTCACCGGCTCCAGATTTAT
CAGCAATAAACCAGCCAGCCGGAAGGGCCGAGCGCAGAAGTGGTCCTGCAACTTT
ATCCGCCTCCATCCAGTCTATCAATTGTTGCCGGGAAGCTAGAGTAAGTAGTTCGC
CAGTTAATAGTTTGCGCAACGTTGTTGCCATTGCTACAGGCATCGTGGTGTCACGC
TCGTCGTTTGGTATGGCTTCATTCAGCTCCGGTTCCCAACGATCAAGGCGAGTTAC
ATGATCCCCCATGTTGTGCAAAAAAGCGGTTAGCTCCTTCGGTCCTCCGATCGTTG
TCAGAAGTAAGTTGGCCGCAGTGTTATCACTCATGGTTATGGCAGCACTGCATAAT
TCTCTTACTGTCATGCCATCCGTAAGATGCTTTTCTGTGACTGGTGAGTACTCAACC
AAGTCATTCTGAGAATAGTGTATGCGGCGACCGAGTTGCTCTTGCCCGGCGTCAAT
ACGGGATAATACCGCGCCACATAGCAGAACTTTAAAAGTGCTCATCATTGGAAAAC
GTTCTTCGGGGCGAAAACTCTCAAGGATCTTACCGCTGTTGAGATCCAGTTCGATG
TAACCCACTCGTGCACCCAACTGATCTTCAGCATCTTTTACTTTCACCAGCGTTTCT
GGGTGAGCAAAAACAGGAAGGCAAAATGCCGCAAAAAAGGGAATAAGGGCGACA
CGGAAATGTTGAATACTCATACTCTTCCTTTTTCAATATTATTGAAGCATTTATCAGG
GTTATTGTCTCATGAGCGGATACATATTTGAATGTATTTAGAAAAATTTTTTAAGGCA
GTTATTGGTGCCGCTTAAACGCCTGGGGTAATGACTCTCTAGCTTGAGGCATCAAA
TAAAACGAAAGGCTCAGTCGAAAGACTGGGCCTTTCGTTTTATCTGTTGTTTGTCG
GTGAACGCTCTCCTGAGTAGGACAAATCCGCCCTCTAGATTACGTGCAGTCGATG
ATAAGCTGTCAAACGGAATTTCGGGCAGCGTTGGGTCCTGGCCACGGGTGCGCC
GGTGTGAAATACCGCACAGATGCGTAAGGAGAAAATACCGCATCAGGCGCTCTTC
CGCTTCCTCGCTCACTGACTCGCTGCGCTCGGTCGTTCGGCTGCGGCGAGCGGT
ATCAGCTCACTCAAAGGCGGTAATACGGTTATCCACAGAATCAGGGGATAACGCA
GGAAAGAACATGTGAGCAAAAGGCCAGCAAAAGGCCAGGAACCGTAAAAAGGCC
GCGTTGCTGGCGTTTTTCCATAGGCTCCGCCCCCCTGACGAGCATCACAAAAATC
GACGCTCAAGTCAGAGGTGGCGAAACCCGACAGGACTATAAAGATACCAGGCGTT
TCCCCCTGGAAGCTCCCTCGTGCGCTCTCCTGTTCCGACCCTGCCGCTTACCGGA

TACCTGTCCGCCTTTCTCCCTTCGGGAAGCGTGGCGCTTTCTCATAGCTCACGCTG
TAGGTATCTCAGTTCGGTGTAGGTCGTTCGCTCCAAGCTGGGCTGTGTGCACGAA
CCCCCCGTTCAGCCCGACCGCTGCGCCTTATCCGGTAACTATCGTCTTGAGTCCA
ACCCGGTAAGACACGACTTATCGCCACTGGCAGCAGCCACTGGTAACAGGATTAG
CAGAGCGAGGTATGTAGGCGGTGCTACAGAGTTCTTGAAGTGGTGGCCTAACTAC
GGCTACACTAGAAGGACAGTATTTGGTATCTGCGCTCTGCTGAAGCCAGTTACCTT
CGGAAAAAGAGTTGGTAGCTCTTGATCCGGCAAACAAACCACCGCTGGTAGCGGT
GGTTTTTTTGTTTGCAAGCAGCAGATTACGCGCAGAAAAAAAGGATCTCAAGAAGA
TCCTTTGATCTTTTCTACGGGGTCTGACGCTCAGTGGAACGAAAACTCACGTTAAG
GGATTTTGGTCATGGAATTAATTCTTAGAAAAACTCATCGAGCATCAAATGAAACTG
CAATTTATTCATATCAGGATTATCAATACCATATTTTTGAAAAAGCCGTTTCTGTAAT
GAAGGAGAAAACTCACCGAGGCAGTTCCATAGGATGGCAAGATCCTGGTATCGGT
CTGCGATTCCGACTCGTCCAACATCAATACAACCTATTAATTTCCCCTCGTCAAAAA
TAAGGTTATCAAGTGAGAAATCACCATGAGTGACGACTGAATCCGGTGAGAATGGC
AAAAGTTTATGCATTTCTTTCCAGACTTGTTCAACAGGCCAGCCATTACGCTCGTCA
TCAAAATCACTCGCATCAACCAAACCGTTATTCATTCGTGATTGCGCCTGAGCGAA
GACGAAATACGCGATCGCTGTTAAAAGGACAATTACAAACAGGAATCGAATGCAAC
CGGCGCAGGAACACTGCCAGCGCATCAACAATATTTTCACCTGAATCAGGATATTC
TTCTAATACCTGGAATGCTGTTTTCCCGGGGATCGCAGTGGTGAGTAACCATGCAT
CATCAGGAGTACGGATAAAATGCTTGATGGTCGGAAGAGGCATAAATTCCGTCAG
CCAGTTTAGTCTGACCATCTCATCTGTAACATCATTGGCAACGCTACCTTTGCCATG
TTTCAGAAACAACTCTGGCGCATCGGGCTTCCCATACAATCGATAGATTGTCGCAC
CTGATTGCCCGACATTATCGCGAGCCCATTTATACCCATATAAATCAGCATCCATGT
TGGAATTTAATCGCGGCCTAGAGCAAGACGTTTCCCGTTGAATATGGCTCATAACA
CCCCTTGTATTACTGTTTATGTAAGCAGACAGTTTTATTGTGTAATCGTTAATCCGC
AAATAACGTAAAAACCCGCTTCGGCGGGTTTTTTTATGGGGGGAGTTTAGGGAAAG
AGCATTTGTCATCATGACCATGACATTAACCTATAAAAATAGGCGTATCACGAGGC
CCTTTCCCTAGGGTCTTCACACTCTATCATTGATAGAGTTAATACGACTCACTATAG
GGTCCCTATCAGTGATAGAGAGAATTCGTACTGAGCACAGCTGTCACCGGATGTG
CTTTCCGGTCTGATGAGTCCGTGAGGACGAAACAGCCTCTACAAATAATTTTGTTTA
AACTAGTTAGATAAGGAGGTTACATATGCACCATCATCACGGTCATCACCACCATC
CGCGTGGTGTTGACCCGAGCCGTAAGGAGAACCACCTGTCTGACGAAGACTTCAA
GGCGGTGTTCGGTATGACCCGTTCTGCGTTCGCGAACCTGCCGCTGTGGAAACAA
CAGAACCTGAAGAAGGAGAAAGGTCTGTTCGGTTCTGGAAGCGCAGCAGCACCTA
AGATGGCATTCACATTGGCCGATCGTGTCACCGAAGAGATGCTGGCAGACAAGGC
AGCCTTGGTCGTGGAGGTAGTTGAGGAGAACTATCACGACGCACCGATTGTTGGA
ATCGCCGTGGTCAATGAACATGGTCGCTTCTTCTTGCGCCCTGAGACTGCGTTGG
CCGACCCACAATTCGTGGCCTGGTTAGGAGATGAAACGAAGAAGAAGTCAATGTT
CGACAGCAAACGCGCAGCCGTAGCTCTGAAGTGGAAAGGAATTGAGCTGTGTGGT
GTGAGTTTCGACCTTCTCTTAGCAGCGTACTTGCTTGATCCCGCTCAAGGCGTCGA
CGACGTGGCAGCCGCTGCCAAGATGAAGCAATATGAAGCGGTGCGTCCGGATGA
GGCTGTGTACGGGAAGGGAGCTAAACGCGCGGTGCCTGATGAACCCGTGCTTGC
TGAGCACTTGGTACGCAAGGCTGCGGCTATCTGGGAGCTGGAGCGTCCCTTCCTG
GATGAGTTGCGTCGCAACGAGCAGGACCGCCTGCTTGTAGAGTTAGAACAGCCTC
TTAGCTCTATTCTTGCCGAGATGGAGTTCGCTGGTGTCAAAGTAGATACCAAGCGC
CTTGAGCAAATGGGTAAGGAGTTGGCTGAACAACTGGGCACAGTGGAACAGCGTA

TCTACGAACTGGCCGGTCAGGAGTTCAACATCAACAGCCCCAAGCAGCTGGGAGT
GATCCTGTTCGAGAAGTTGCAGCTGCCAGTATTGAAGAAGACTAAGACTGGCTACA
GTACCTCGGCTGACGTACTGGAGAAGCTGGCTCCTTACCATGAGATCGTGGAGAA
CATCTTGCACTACCGCCAGCTGGGCAAGCTGCAGTCTACCTACATTGAGGGTCTG
TTAAAGGTCGTGCGTCCAGACACGAAGAAGGTGCATACGATCTTCAATCAGGCGC
TGACCCAAACTGGTCGTTTGTCGTCCACAGAGCCCAATCTTCAGAATATCCCTATT
CGTCTTGAGGAAGGCCGCAAGATTCGCCAGGCCTTCGTTCCTTCGGAATCGGACT
GGCTGATCTTCGCAGCAGATTACTCACAGATCGAGCTTCGCGTGTTGGCACATATC
GCGGAGGATGACAACTTAATGGAGGCGTTCCGCCGCGATCTGGATATCCATACTA
AGACCGCGATGGATATCTTCCAAGTGTCAGAAGACGAGGTAACACCGAACATGCG
ACGCCAGGCGAAAGCGGTTAACTTCGGCATCGTCTACGGCATCAGCGACTATGGC
CTGGCCCAGAACTTGAACATCAGCCGCAAGGAGGCAGCCGAGTTCATCGAGCGCT
ACTTCGAGAGTTTCCCAGGTGTGAAGCGTTATATGGAGAATATCGTACAAGAGGCG
AAGCAGAAAGGCTACGTGACCACGCTGTTACACCGTCGTCGCTACCTTCCTGATAT
CACTAGCCGTAACTTCAATGTACGTTCCTTCGCCGAACGCATGGCGATGAATACCC
CCATCCAGGGGTCAGCTGCAGATATCATCAAGAAAGCTATGATCGACTTAAACGCT
CGTCTGAAGGAAGAACGCTTACAGGCGCACCTCTTACTGCAAGTCCATGACGAATT
GATCCTTGAGGCGCCCAAGGAAGAGATGGAGCGTCTTTGCCGTCTGGTGCCGGA
AGTAATGGAACAGGCCGTCACGCTGCGCGTACCTCTGAAAGTCGATTACCACTAC
GGCTCCACCTGGTATGACGCCAAGTAAGG
